# The role of effort in adaptation to split-belt locomotion

**DOI:** 10.1101/2025.05.07.652669

**Authors:** Rachel M. Marbaker, Yolande Idoine, Alaa A. Ahmed

## Abstract

In many motor learning tasks, the process of *error* reduction is mirrored in a process of *effort* reduction, where metabolic cost and or muscle co-contraction decrease gradually in tandem with error. Effort reduction may be incidental to the learning process, but high error movements can be more effortful and a drive to minimize effort costs could also play a role in motor learning. In this study, we focused on the effort requirements of the task, asking whether task (background) effort cost affects learning, aftereffects, or relearning in a split-belt walking task. We hypothesized that greater effort costs would amplify the need to reduce effort and accelerate learning. Alternatively, we hypothesized that greater effort requirement could compromise and slow the learning process. Participants in high, low, and control effort groups completed a split-belt walking task while wearing a weighted vest with 15% body mass, 5% body mass, or the vest only, and we assessed step length asymmetry throughout the protocol. Step length asymmetry changed similarly between groups, with similar rates and extent of learning and relearning and similar patterns of aftereffects when the split-belt perturbation was removed. We found no significant effect of task effort on the process of split-belt adaptation, suggesting that the brain’s response to gait asymmetry and ability to adapt to novel dynamics is relatively unchanged by background effort requirements of the task.

**NEW & NOTEWORTHY:** Despite the brain’s sensitivity to effort cost and willingness to adjust gait parameters to minimize cost, the process of split-belt adaptation was unaffected by increased effort requirements. This finding provides a foundation for further research into performance-dependent effort cost. Additionally, modest increases in effort should be further explored in rehabilitation applications where higher effort requirements may help build strength and fitness without impairing motor learning.

## INTRODUCTION

The brain ably learns and adapts to novel environments and perturbations, coordinating sensing, action selection, motor response, and error correction to flexibly respond to a range of internal (pain, muscular exhaustion, energetic cost) and external (reward, punishment, salience) information. The brain’s process of motor learning is characterized by gradual error reduction, so that when a person’s movement is disrupted by an internal or external change to the environment, they respond by adjusting their motor strategy (explicit learning) or by retraining an internal map between motor commands and outcomes (implicit learning). These two processes contribute together to compensate for the change, reducing error and improving performance. Motor learning is sensitive to external factors including reward and punishment (1, 2), while internal factors like pain and fatigue may change patterns of muscle recruitment (3, 4). Could effort play a role in learning similar to these factors? Effort, the physical cost of completing a movement, is minimized in movement selection, both consciously in decision-making tasks and subconsciously where people walk and run at effort-minimizing speeds and gait characteristics (5–7). Furthermore, in novel effort landscapes, individuals adjust movements and strategies to find new minima (8–11). Thus, effort, like reward and punishment, is a factor in the trade-off between movement options (12). The factors that influence movement selection notably overlap with the factors that influence learning. For example, reward has been shown to increase the vigor of reaching movements and also to increase learning rate in visuomotor rotation (2, 13).

Effort can also offer a window into the learning process. Parallel to error reduction during learning, there is a pattern of effort reduction. That is, effort costs decay as performance improves, and this finding holds true in a variety of task paradigms. In reaching tasks perturbed by force field or visuomotor rotation, changes in effort cost are observed in net metabolic cost and muscular co-contraction, both of which are elevated early in learning when performance error is high (14, 15). Co-contraction and net metabolic cost decay over the course of the learning block, matching the gradual reduction of error as participants become more adept at accommodating the perturbation (14, 15). Similarly, in walking, while participants adapt to split-belt treadmill locomotion, metabolic costs are initially high and decay gradually over the course of learning split-belt gait (16).

In split-belt walking a typical adaptation period lasts between 8 and 15 minutes, during which step length asymmetry shifts from far longer steps on the slow leg to nearly symmetric step lengths, though usually with some residual bias toward longer steps on the slow leg (17–20). In a computational model of split-belt walking, the optimal gait pattern allowed the participant to use the net positive work of the treadmill, reducing work from the muscles (9). The optimal gait was *asymmetric*, biased toward longer steps on the fast leg, contrasting the bias most often observed in late adaptation. (9). With prolonged split-belt exposure or specific training, people adjusted their gait to take advantage of the belt-speeds, suggesting that participants may be sensitive to the available effort savings during extended exposures to perturbed effort landscapes (8, 9).

Factors like perturbation magnitude (belt-speed ratio or difference), attention, varied repetition schemes, and anterior-posterior ground reaction forces have been implicated as mediators of adaptation and retention (18, 21, 22, 25–31). Of these mediators, adaptation to belt-speed differences and modulated ground reaction forces can be most readily interpreted in the context of effort cost. For various belt-speed combinations and ratios, Butterfield and Colins (2022) showed that the metabolic cost of split-belt walking is overall similar to the metabolic cost to tied-belt walking at the average speed of the two belts, though larger belt-speed differences also incur slightly higher cost (32). In learning, larger belt speed ratios resulted in further extent of learning (more progress toward symmetry) and faster *re*learning (22). Possibly, the higher speed ratios engage more explicit strategy, but an alternative hypothesis is that the higher belt speed ratio elevated cost of asymmetry, pushing participants toward more efficient and more symmetric gaits (22). Kinetic and muscle activity characterizations of split-belt showed that anterior ground reaction forces (braking) had the most profound changes during learning (27). However, in a study where participants adapted to split-belt on declined, level, or inclined belts, it was participants in the inclined treadmill group, with sapped braking forces and augmented propulsive forces, who learned faster and to a greater extent (29). In this case, improved learning could be attributed to either the greater propulsive force demands or to higher background effort cost (29). In both the belt-speed ratio studies and the ground reaction force studies, an effect of background effort cannot be detangled from other characteristic gait changes the experimental manipulations. Here, we try to disassociate the effect of background effort on split-belt adaptation by modulating the amount of additional mass carried in a vest during the task protocol.

The addition of modest weight has several factors that make it a useful manipulation for background effort. The weighted vest increases the cost of walking independent of any split-belt performance or symmetry-related outcome in a way that is familiar from day to day life (for example, carrying a backpack), and with little effect on gait kinematics (33, 34). While the added mass does increase ground reaction forces, the change is uniform across directions and importantly requires reciprocal changes in braking and propulsion. Based on these factors and other literature on effort and learning, we predicted three potential outcomes for the role of background effort during split-belt adaptation.

Our first prediction was that additional background effort, by uniformly augmenting effort cost across the task, will *upregulate* effort cost during effort during the early, high-error phase of learning. With an upregulated effort cost, there may be greater urgency reach a low-error-low-effort state. That is, the higher the magnitude of effort, the more urgently the brain might search for a more efficient movement. In this case, we expected that the higher background effort costs would drive faster adaptation (Fig. 1A). Several potential mechanisms could accelerate learning: i) increased error sensitivity if, for example, effort cost increases movement variability, encouraging the brain to weigh and attend to different error signals (35); ii) the relative magnitude of the error signal could increase with background effort if, for example, the proprioceptive signal, known to inform adaptation to incline walking, increases alongside the muscular demand; and iii) in perhaps the simplest mechanism for accelerating learning, the additional effort cost may augment the perception of belt-speed differences, which may influence the strategy of adaptation (36–38).

**Figure 1:**
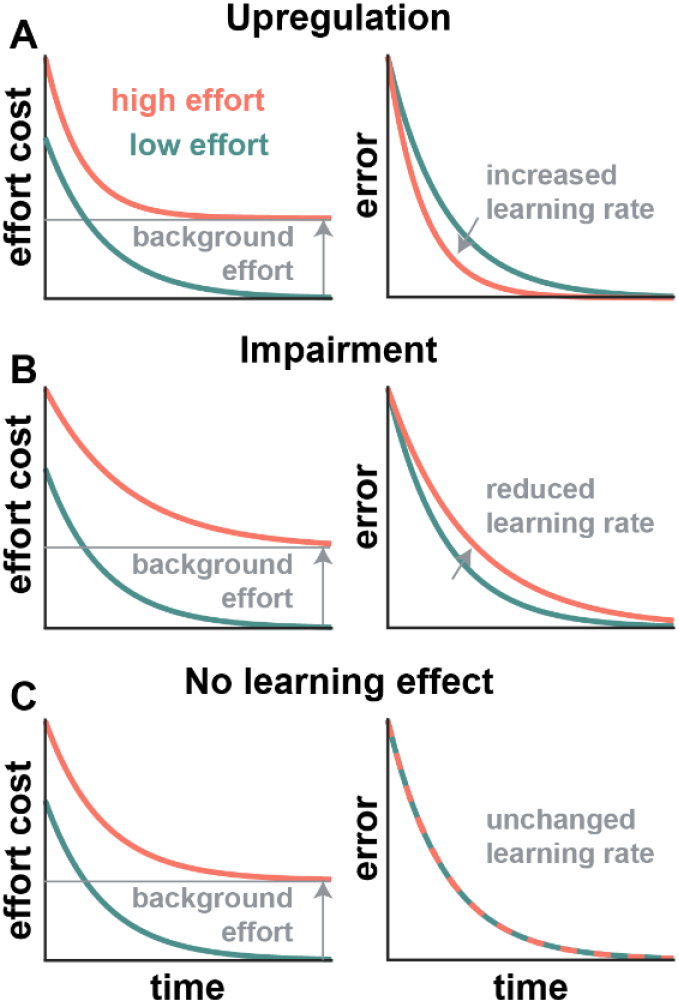
Schematic for different relationships between effort and error over time in a learning task illustrated for A) upregulation where effort makes error reduction more urgent, B) impairment where effort slows learning by distracting or discouraging error reduction, and C) indifference where effort has no effect on error reduction.

In an alternative prediction, background effort could *impair* motor learning by distracting the learner or by quelling motivation to adapt by reducing the quality of the learning environment. Distraction with a secondary task slowed split-belt adaptation (18, 39). While the additional mass is probably not cognitively distracting, any discomfort or pain associated with walking asymmetrically or carrying extra mass could distract the learner. Previous work has shown that pain influenced muscle recruitment during learning and impaired retention in arm reaching and locomotor adaptation (3, 40–43). Additional mass could also change how a learner responds to and recalls the environment. While the absence of reward- or performance-based motivation during split-belt adaptation dilutes any clear reward rate, the presence of greater effort cost could discount the average “goodness” of the learning environment. Environmental goodness has been associated tonic dopamine levels, which influence motivation and overall engagement in the task (44). Thus, if the added mass reduces the quality of the environment, it may impair learning by diminishing engagement (Fig. 1B).

Finally, it is also possible that motor learning and recall processes are *indifferent* to the background effort cost (Fig. 1C). This could be because error and asymmetry take precedence over the effort cost especially in relatively short exposures where there is little accumulation of fatigue or discomfort. The perceived risk related to walking on a treadmill and the unfamiliarity of the split-belt condition may substantially outweigh any signal related to a modest increase in effort. In this case, we would expect that changing the background effort will not affect the learning process. The addition of effort could change muscle activation and gait control strategies without changing the rate or extent of the adaptation process, similar to observations on the effect of pain (3, 40, 43).

To test the effect of background effort on split-belt adaptation, we measured the step length asymmetry for three different effort condition groups over two 10-minute split-belt adaptation blocks which were separated by a 15-minute washout block. Each effort condition included 21 naive participants wearing a weighted vest (15% body mass, 5% body mass, and vest only for the high, low, and control conditions respectively). Despite the important role of effort in movement selection and the clear trend of effort reduction over the course of learning, we found no evidence of a causal or modifying role of background effort costs on adaptation to a split-belt perturbation.

## METHODS

### Subjects

Participants in the split-belt walking study (n = 66; 36W, 30M) with an average age of 24.5 ± 4.22 (mean ± standard deviation) years all signed a consent form approved by the Institutional Review Board at the University of Colorado Boulder. Participants were healthy and reported no known neurological or balance disorders and had no recent injuries affecting the lower limb. All participants were compensated for their time. Participants were evenly divided into randomly assigned effort conditions—high, low, and control—which were differentiated by the additional mass worn in a vest during the task. Three participants were excluded because they were unable to walk comfortably on the treadmill or unable to let go of the handrails during fast tied-belt or split-belt blocks. Ultimately, the three groups (low, high, and control) consisted of 21 participants each. Of the participants in the low or high effort conditions, 22 returned for a second visit within 14 days to repeat the task in the opposite condition (11 participants high-low and 11 participants low-high, see Fig 2C).

**Figure 2:**
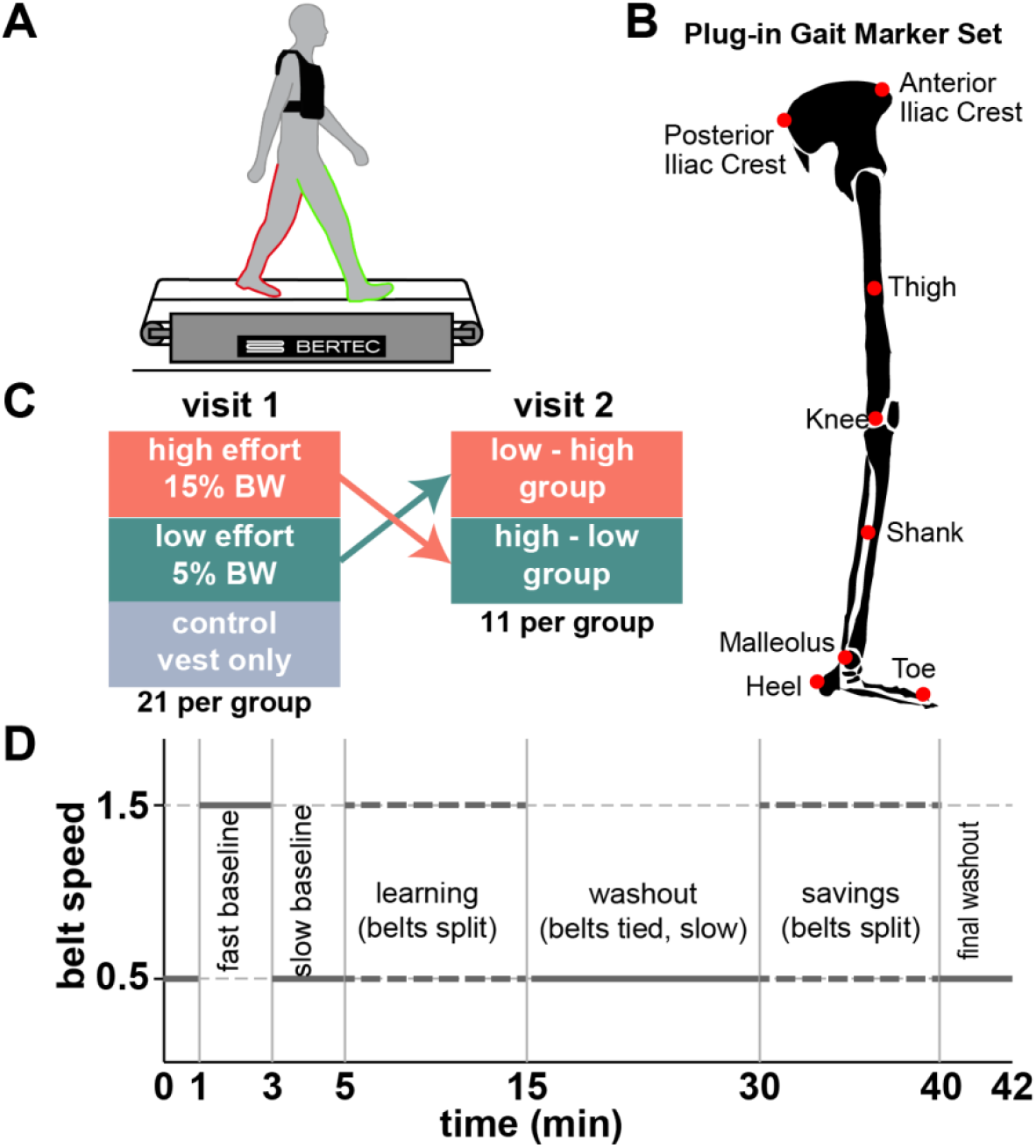
Methods. A) a schematic of split-belt walking showing the participant wearing the weighted vest and each foot on a separate belt. B) Plug-In Gait marker set showing the eight markers on the right leg. Participants who had markers on the feet-only had the toe, heel, and malleolus markers on each foot. C) Effort condition groups in the first and second visit. D) The 42-minute split-belt protocol with tied-belt baseline blocks, followed by the first split-belt exposure, a tied-belt washout block, a second split-belt exposure, and a final tied-belt washout.

### Experimental setup

All participants walked on a treadmill with a separate belt for each foot (Bertec Instrumented Treadmill). They wore a weighted vest for the duration of the task with added mass determined by their effort group: no additional mass (control condition), 5% of body mass (low effort condition), or 15 % of body mass (high effort condition). Mass was added to the vest in 1.13 kg (2.5 lb) increments, so the percentage of body mass was rounded down from 5% body mass in the low effort condition and rounded up from 15% body mass in the high effort condition. Treadmill belt speeds were always 0.5 m/s (slow speed) or 1.5 m/s (fast speed) and accelerated from a stop at a rate of 0.5 m/s. The treadmill belts accelerated from or to a stop and never changed speed while moving. During tied-belt blocks, the two belts moved at the same speed. During split-belt blocks, one belt moved at the fast speed while the other moved at the slow speed. Participants were randomly assigned a “fast leg,” right or left, and the assignments were kept even within groups. For participants who visited the lab twice and completed the task in high effort and low effort conditions, the order of conditions was counterbalanced while participants’ “fast leg” was consistent between visits so that any observed changes in learning would not be confounded with differences in generalization between legs.

### Protocol

Participants completed seven walking blocks with short (less than 30 s) pauses between blocks. Participants were warned of stops or starts by a series of three beeps and instructed to prepare to grab the handrails if needed, but let go within 5 seconds of the trial starting to minimize effects of handrail grasping on the adaptation process (45). The first block was a baseline block and consisted of 1 minute walking with both belts at slow speed (tied-belts). This was included to help accustom participants to walking on the two separate belts. The second and third blocks were two minutes with tied-belts at the fast and slow speed respectively to record baseline gait parameters during symmetrical walking. During the subsequent learning block, the split-belt perturbation (belt ratio 3:1 with speeds at 1.5 m/s and 0.5 m/s) was introduced and remained constant for 10 minutes while participants walked continuously. Following learning, adaptation was washed out during a 15-minute block of continuous walking with tied-belts at the slow speed (washout block). During relearning (savings block), the belt speeds split again for 10 minutes, keeping fast leg consistent with the first learning block. Participants finished the task with 2 minutes walking with tied-belts at the slow speed to ensure a safe return to overground walking. The subset of participants who walked in both effort conditions returned within two weeks and completed the same seven block walking task while wearing the weighted vest for the opposite effort condition. We denote these groups based on the order of their visits as low-high or high-low.

### Data collection

During each block of the walking task, ground reaction forces and moments were collected via force plates embedded in the treadmill belts at a sampling rate of 1000Hz. For 45 participants, we collected motion capture data at a frequency of 100 Hz using the Plug-In Gait marker set model with 16 reflective markers placed on the lower body (Vicon Vero Cameras; Vicon Nexus software). The Plug-In Gait Model (Fig 2B), includes markers on the right and left toe, heel, malleolus, knee, thigh, and anterior and posterior superior iliac crests. For the remaining 18 participants, we collected motion capture data for only the feet with markers on the right and left toe, malleolus, and heel. Data was low pass filtered and processed manually to fill gaps using Vicon Nexus software and analyzed using custom MATLAB scripts.

### Metrics

Our metrics for this study are hierarchical, consisting of step, stride, and adaptation measures. We defined our step metrics based on gait events and phases: stance refers to the phase of the gait where the foot is on the ground, starting with heel strike and finishing with toe off; swing refers to the movement between toe off and heel strike when the foot is in the air; and a step corresponds to the stance and swing for a single leg. To identify gait events, we used vertical ground reaction force (100 N threshold) to identify heel strike and toe off gait events using custom processing scripts in MATLAB.

Kinematic step metrics included: step length (SL), the distance between malleolus markers at the time of the forward foot heel strike; step time, the time between heel strikes on opposite feet; and step width, the frontal plane distance between the heel markers at the time of the forward foot heel strike (6). We also compared step length, time, and width variability between groups. We quantified variability as variance. Higher variance has been implicated in faster learning and effort optimization, potentially because it increases “exploration” error and effort landscapes (10, 46). As the traditional learning metric, step length asymmetry exploration may be a strategy to accelerate learning. Previous work has suggested that effort optimization in split-belt walking may correspond to a step time asymmetry optimum, so step time variance may also help drive faster adaptation by exploring the effort landscape (24). Step width variability is a measure of balance and allowed us to quantify, in baseline, the effects of adding weight above the center of mass and, in learning, strategies for maintaining or improving asymmetry (47, 48). Kinetic step metrics included: peak braking force, peak propulsive force, and peak vertical ground reaction force. We also calculated propulsive impulse (time integrated propulsive force in the stance phase). We included propulsive impulse in addition to peak propulsive force because it incorporates stance duration. All kinetic metrics were normalized to body mass.

We used stride metrics to examine the relationship between steps on the fast and slow legs. A stride lasts from the heel strike on one foot until the subsequent heel strike on the same foot. We selected our stride metrics based on previous work focusing on asymmetry measures. The most common, step length asymmetry (SLA) shows robust patterns of adaptation and savings (20, 21, 24, 49). We defined step length asymmetry as

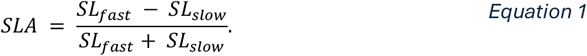

It describes the ratio of step lengths between the leg on the fast belt and the leg on the slow belt. Over the course of learning, it gradually adjusts toward symmetry (SLA = 0). Step time and step width asymmetries were defined using the same construction: fast – slow / (fast + slow). Each stride metric is a measure of asymmetry for each stride in the protocol block.

Plotted against stride number, stride metrics like step length asymmetry describe the learning process. To match common split-belt analyses in the literature, we compared step length asymmetry during blocks of strides, representing different phases of the learning process (initial: strides 1-5, early: strides 6-30, late: strides 31-200, and plateau: final 30 strides) (21, 22, 28).

We characterized the learning process as a whole using adaptation metrics: step length asymmetry, strides to plateau and parameters derived from state space and exponential model fits. Strides to plateau is a curve-agnostic count of the number of strides to reach steady state asymmetry (plateau) during learning, washout, and relearning blocks. Based on previous work, we defined the threshold for plateau based on the mean and standard deviation for the last 30 strides of the block. We identified the first stride that was followed by five consecutive strides falling inside this plateau range as the strides-to-plateau value for each participant (18).

We also fitted a descriptive exponential model to each participants’ step length asymmetry curves in learning, washout, and relearning.. We calculated fit parameters according to an exponential function: *SLA*(*t*) = *Ae*^*βt*^ + *c*. In the fit function, the exponential coefficient represented the change in asymmetry (Δ*SLA*), the exponential rate (β) represented the learning rate, and the exponential constant represents the final asymmetry or plateau (*SLA*_*final*_). We fit the exponential function in a custom script using MATLAB’s *fminsearch* function.

Lastly, we fit the learning parameters of a state space model. The state space learning model is recursive, predicting an updated gait plan (*x*_*n*_) for each stride *n* based on the previous gait plan and the error from the previous stride action *y*(*n*). The error of the gait action is calculated as the difference between the gait action and the split-belt perturbation: *e*(*n*) = *p*_*n*_ − *y*(*n*). Then, the gait action is assumed to be equivalent to the gait plan (*y*(*n*) = *x*_*n*_) and the update model takes the form:

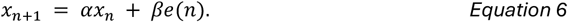

In the update, the gait plan from the previous trial is weighted by a remembering factor, α, describing how much of the prior gait plan is remembered and the error from previous trial is weighted by a learning rate, β, describing the participant’s error sensitivity. We fit both remembering factor, α, and learning rate, β, in the interval [0,1] with a custom script using MATLAB’s *fmincon* function.

### Statistics

We used the baseline blocks to quantify the effects of effort condition on gait measures. We examined the effect of belt speed and effort condition on step kinetics and kinematics (step length, step time, step width, peak braking, peak propulsion, vertical ground reaction force, and propulsive impulse) using repeated measures ANOVAs with baseline block belt speed (within-subjects) and effort condition (between-subjects) as factors.

#### Learning

Our primary measure of learning is step length asymmetry. Step length asymmetry was compared between groups in each phase of the adaptation blocks (learning, washout, and relearning). The phases refer to sets of trials whose average can represent different parts of adaptation process. Adaptation (or de-adaptation) blocks are divided into initial (strides 1-5), early (strides 6-30), late (strides 31-200), and plateau phases (final 30 strides) (21, 22, 50). We compared groups’ symmetry across all four phases using mixed repeated measures ANOVAs with effort condition (between-groups) x learning phase (within-group). To assess savings, we created a larger three-way ANOVA that included symmetry phase data for both learning and relearning blocks (effort condition x block (learning or relearning) x phase). Strides to plateau was compared using a one-way anova for each block, and mixed anova to quantify savings.

Step kinematics, kinetics, and variability were compared using three-way repeated measures ANOVAs that included effort condition, phase and an additional variable for the fast leg or slow leg (within-group). We used estimated marginal means with a Bonferroni correction to unpack RM-ANOVA interactions as needed.

We compared the state space learning parameters and exponential fits between groups using one-way ANOVAs for each block and mixed ANOVA to quantify savings between learning and relearning. Fit parameter comparisons were similar when the data was fit across all strides of the learning block to when the data was fit using only the first 200 strides, so the state space parameters presented were fitted to all strides in the learning block. We used independent t-tests for planned comparisons between group pairs.

During the second visit, we used the same repeated measures ANOVA structure to compare effort conditions (high-low and low-high) over the adaptation block phases. We used independent t-tests to compare fit parameters between the second visit groups.

We assigned numbers to each exposure to the split-belt perturbation with 1 and 2 identifying learning and relearning during the first visit and 3 and 4 identifying learning and relearning during the second visit. We were interested in the rate and extent of learning across the four exposures and so focused our analysis on learning rates from the state space and exponential fits and on the asymptote of step length asymmetry. We used a three-way ANOVA across all four exposures (learning and relearning on the first and second visit) and all phases (initial, early, late, and plateau) to compare between the high-low and low-high groups.

We collected data from a sample size consistent with comparable published studies. Expecting that effort might affect split-belt locomotion in a manner similar to distraction or pain, we used locomotion data published by Malone and Bastian (2010) on the effect of distraction and by Galgiani et al. (2024) on the effect of pain to conduct our power analysis. For similar effect sizes, at 0.95 power and 0.05 significance level, we determined a sample size of 21 participants per group.

## RESULTS

Comparing adaptation to split-belt walking between three effort condition groups of naïve participants, we found no evidence of differences in learning, aftereffects, or savings among our step, stride, and adaptation measures. The subset of individuals in the high and low effort groups that returned to the lab for a second visit showed strong savings with no difference between groups related to the order of effort conditions.

### Participants across effort conditions walked with similar and symmetrical gait during baseline

As a measure of baseline walking performance, all participants completed 2-minute blocks of symmetrical walking on the treadmill at the fast speed (1.5 m/s) and at the slow speed (0.5 m/s) while wearing the weighted vest. Before examining the learning phases, we wanted to compare baseline performance between the groups. Steps in fast baseline were longer than those in slow baseline and similar between effort groups (RM-ANOVA comparing fast and slow baseline between effort conditions; main effect of speed (fast or slow baseline): p = 4.24e-48; no main effect of effort condition: p = 0.924; interaction p = 0.589). Similarly, step times were shorter in fast baseline (main effect of speed: p = 8.33e-38) and similar between effort conditions (p = 0.741, interaction p = 0.436). Step widths were similar between fast and slow baseline and between effort conditions suggesting that participants did not accommodate the added weight with wider steps in baseline, nor did participants respond to faster treadmill speeds by widening their gait (RM-ANOVA, speed p = 0.626, effort condition p = 0.553, interaction p = 0.247).

Step length variance in fast and slow baseline was similar between the high, low, and control effort groups (repeated measures ANOVA no main effects of effort condition F(2,56) = 1.11, p = 0.319 or speed F(1,56) = 2.04, p = 0.158). Step length variance was sensitive to both effort condition and speed (main effect of effort, F(2,56) = 3.40, p = 0.039 and speed, F(1,56) = 2.04, p = 6.12e-04). The interaction was also significant (F(2,56) = 3.45, p = 0.039) suggesting that the directional effect of speed and step time variance was mixed between the effort groups. Step width variance was similar between groups (RM-ANOVA no main effect of effort condition, F(2,56) = 0.579, p = 0.564, or block, F(1,56) = 2.23, p = 0.053, no interaction F(2,56) = 0.623, p = 0.623). The effect of speed trended toward significance based on high step width variance at higher speeds. Lastly, we verified that step lengths in fast and slow baseline were symmetric by checking comparing the step length asymmetry for last 30 strides in the two baseline conditions to full symmetry (asymmetry = 0) using t-tests (all p values > 0.1).

In summary, while walking speed altered step length and step width variability, gait kinematics when the belts are tied were not influenced by the added mass.

### Added mass increased braking and propulsive forces

Since prior work has implicated ground reaction forces as mediators of split-belt adaptation (27, 29), we compared ground reaction profiles between effort groups, looking first at peak braking and propulsion forces as well as propulsion impulse (Figure 3).

**Figure 3:**
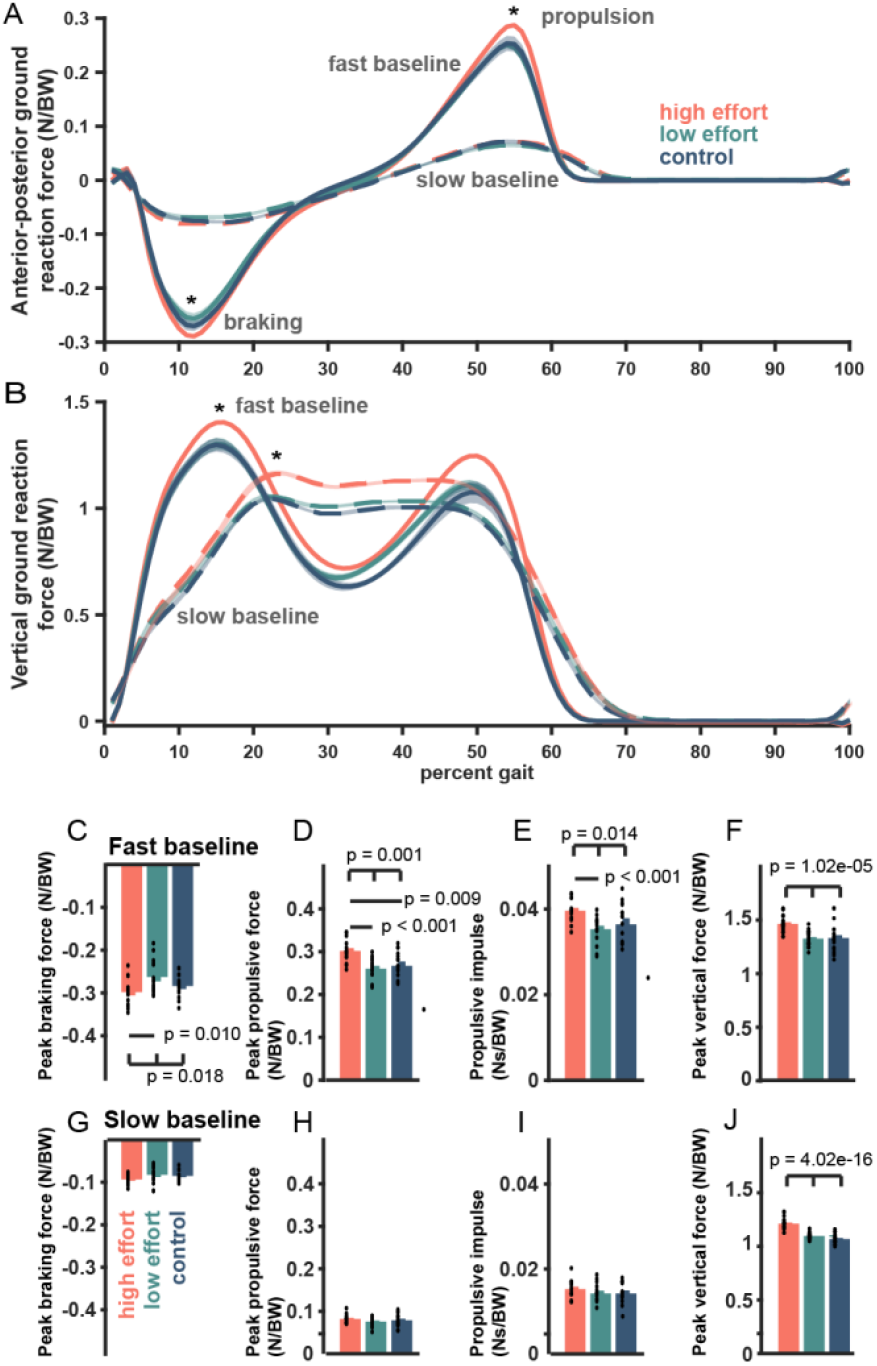
Baseline ground reaction force. A) normalized anterior-posterior and B) vertical ground reaction forces for fast (solid) and slow (dashed) baseline vs percent gait cycle. Traces are subject averages. Shaded regions are 1SEM. Bar plots present mean values of peak braking, peak propulsion, propulsive impulse, and peak vertical force for fast baseline(C – F) and slow baseline in (G – J). Dots represent individual participants.

Peak propulsive force was higher in fast baseline and for the high effort group (RM-ANOVA main effect of effort condition, F(2,60) = 8.97, p = 3.91e-04, and speed, F(1, 60) = 3869, p = 3.37e-56, with significant interaction, F(2,60) = 12.36, p = 3.19e-05, Figure 3A,D,H). Accordingly, peak braking forces were higher in fast baseline and for the high effort group between effort conditions and speed (RM-ANOVA main effect of effort condition, F(2,60) = 5.61, p = 0.006; and speed, F(1,60) = 3091, p = 2.54e-53: and interaction, F(2, 60) = 5.66, p = 0.006) with higher braking in fast baseline walking and in the high effort group (Fig 3A,C,G).

Vertical ground reaction force was higher in fast baseline and for the high effort group consistent with our weight-based effort intervention (RM-ANOVA main effect of effort condition, F(2, 60) = 34.3, p = 1.18e-10; and speed, F(1,60) = 504, p = 6.96e-36; and no interaction F(2,60) = 0.882, p = 0.419; Figure 3B).

Propulsive impulse was higher in fast baseline, but similar between effort conditions (RM-ANOVA main effect of speed, F(1,60) = 412.2, p = 8.19e-29, but not of effort condition F(2, 60) = 2.25, p = 0.114; and no interaction F(2,60) = 1.76, p = 0.181). Possible trending significance for an influence of effort condition was driven by the high effort group in fast baseline, Figure 3A, E, I).

Overall, the added mass increased propulsive, braking, and vertical ground reaction forces and propulsive impulse, particularly in the fast baseline block.

### During learning, all groups gradually shifted from braking with the slow leg to equal braking and propulsion

Next, we turned to the learning block. Participants in all three effort groups adjusted braking and propulsion forces over the course of split-belt exposure. For the first few steps of the split-belt exposure, participants braked with the slow leg and pushed off with the fast leg (Figure 4A, left). As learning progressed, slow leg propulsion and fast leg breaking increased. By final learning, braking and propulsion forces were balanced within each leg. Vertical ground reaction forces (Figure 4B) were higher on the slow leg than on the fast leg throughout learning. By the end of learning, the fast leg vertical ground reaction force shifted from a single peak to the double peak characteristic of rolling from heel-to-toe during stance.

**Figure 4:**
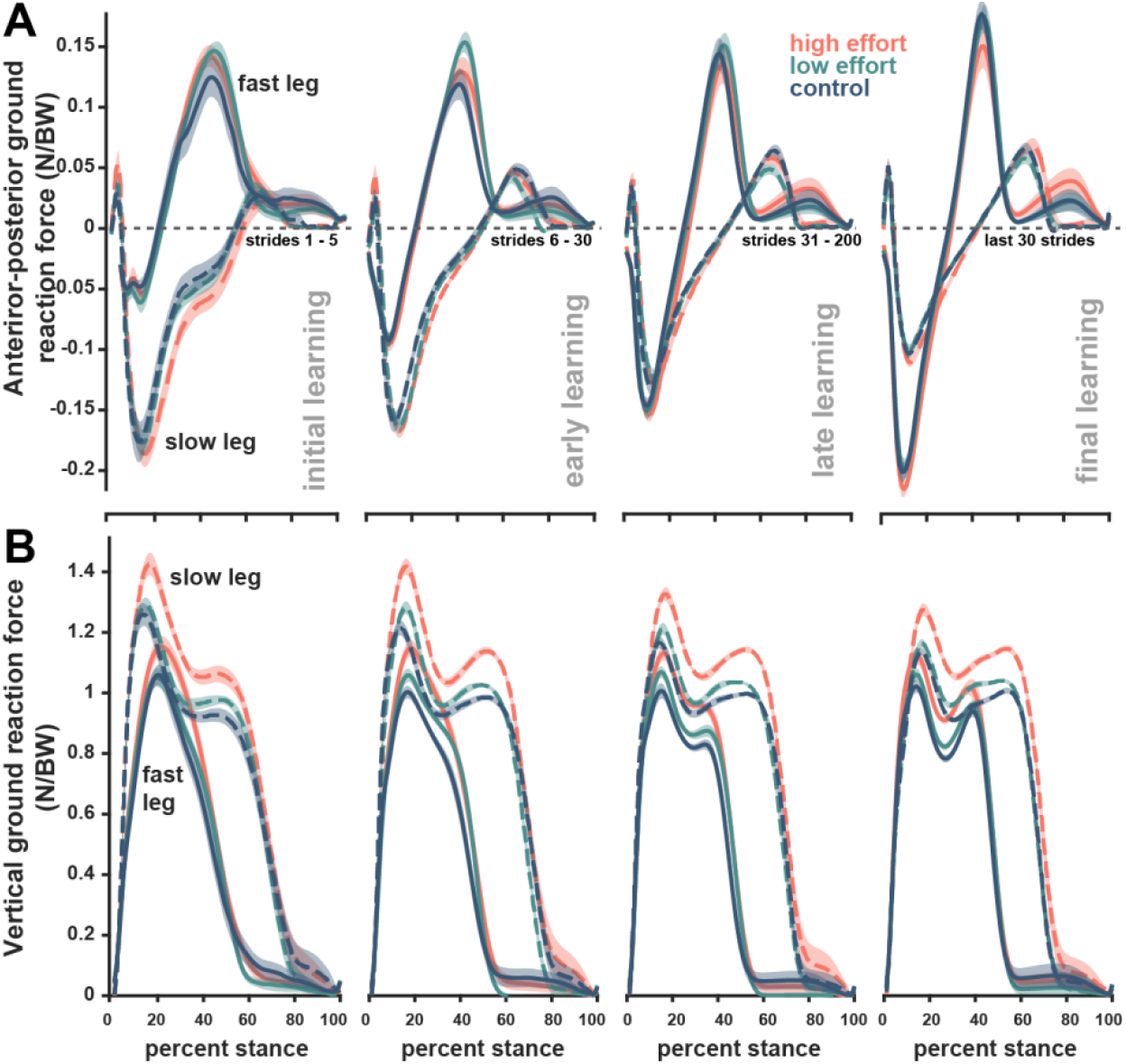
Learning ground reaction forces. (A) anterior-posterior (braking and propulsive) ground reaction forces. (B) vertical ground reaction forces. Traces are averages over binned strides during initial, early, late, and final learning. Dashed lines indicate the slow leg while solid lines indicate the fast leg. Shading is ±1SEM.

Ground reaction force asymmetries represent another potential metric of learning as both propulsion and braking have been implicated as mediators of split-belt adaptation (19, 27, 50). Of the ground reaction force asymmetries we assessed, none were sensitive to effort condition (vertical ground reaction force asymmetry, F(2,60) = 0.314, p = 0.732; peak propulsive force asymmetry, F(2, 60) = 0.966, p = 0.386; peak braking asymmetry F(2, 60) = 0.158, p = 0.854; impulse asymmetry, F(2, 60) = 0.933, p = 0.399), though some changed over the course of learning. Consistent with previous findings, propulsive force asymmetry was unaffected by learning phase (F(2.12, 127.3) = 1.55, p = 0.214; nonsignificant interaction F(4.24, 127.3) = 0.751, p = 0.567) (27). Peak vertical force asymmetry was initially negative and increased toward symmetry (phase F(1.87, 112.31) = 59.63, p = 1.14e-17; interaction F(3.74, 112.3) = 0.524, p=0.702), while impulse asymmetry was initially high (and positive) and decreased toward symmetry (F(2.08,125) = 27.1, p = 9.01e-11, interaction F(4.17,125) = 1.46, p = 0.217).

Taken together, both the magnitude and asymmetry of ground reaction forces adapted over the course of learning, with no significant effect of effort condition.

### Effort condition did not influence adaptation of step length asymmetry

Over the course of exposure to the split-belt, the step length asymmetry progressed from asymmetry toward symmetry. For strides early in the block, step length asymmetry is negative for all groups indicating that participants responded to the split-belt perturbation by taking longer steps on the slow leg. Over time, participants in all groups adjusted their relative fast and slow step lengths to be more similar, although a slight bias toward longer steps on the slow leg persisted for the duration of the 10-minute learning block.

Consistent with gradual learning, step length asymmetry changed between phases of learning (main effect of phase, F(1.99, 103.7) = 260,p = 0.4.42e-41). However, there was no effect of effort condition or interaction between effort condition and learning phase (effort condition, F(2,52) = 0.261, p = 0.771, interaction, F(3.99, 103.7), p = 0.951).

None of the groups achieved their slow baseline symmetry over the course of the learning block (paired t-tests between learning plateau and slow baseline step length asymmetry: high t(20) = -6.31, p = 3.67e-06; low t(19) = -3.76, p = 0.001; control t(19) = -5.80, p = 1.37e-05). For thoroughness, we also compared plateau learning against perfect symmetry (step length asymmetry equal to 0): none of the groups achieved symmetry during adaptation (high t(20) = -3.76, p = 0.001; low t(19) = -3.14, p = 0.005; control t(20) = -4.18, p = 4.26e-04).

There was also no effect of added mass on strides to plateau. For initial learning, the first stride to the plateau was 128 ± 16.90 strides for the high effort condition, 170 ± 17.17 strides for the low effort condition, and135 ± 15.49 strides for the control condition (One-way ANOVA: F(2,53) = 1.77, p = 0.181).

### Model-based fits

From the state space model, remembering factor, learning rate, and initial error were similar between groups (all p>0.30, Figure 5C-F). The exponential coefficient represents the change in asymmetry (Δ*SLA*), the exponential rate (β) represents the learning rate, and the exponential constant represents the final asymmetry or plateau (*SLA*_*final*_). One-way ANOVAs comparing the fit parameters between groups revealed no effect of effort condition (all p > 0.07, Figure 5G-J).

**Figure 5:**
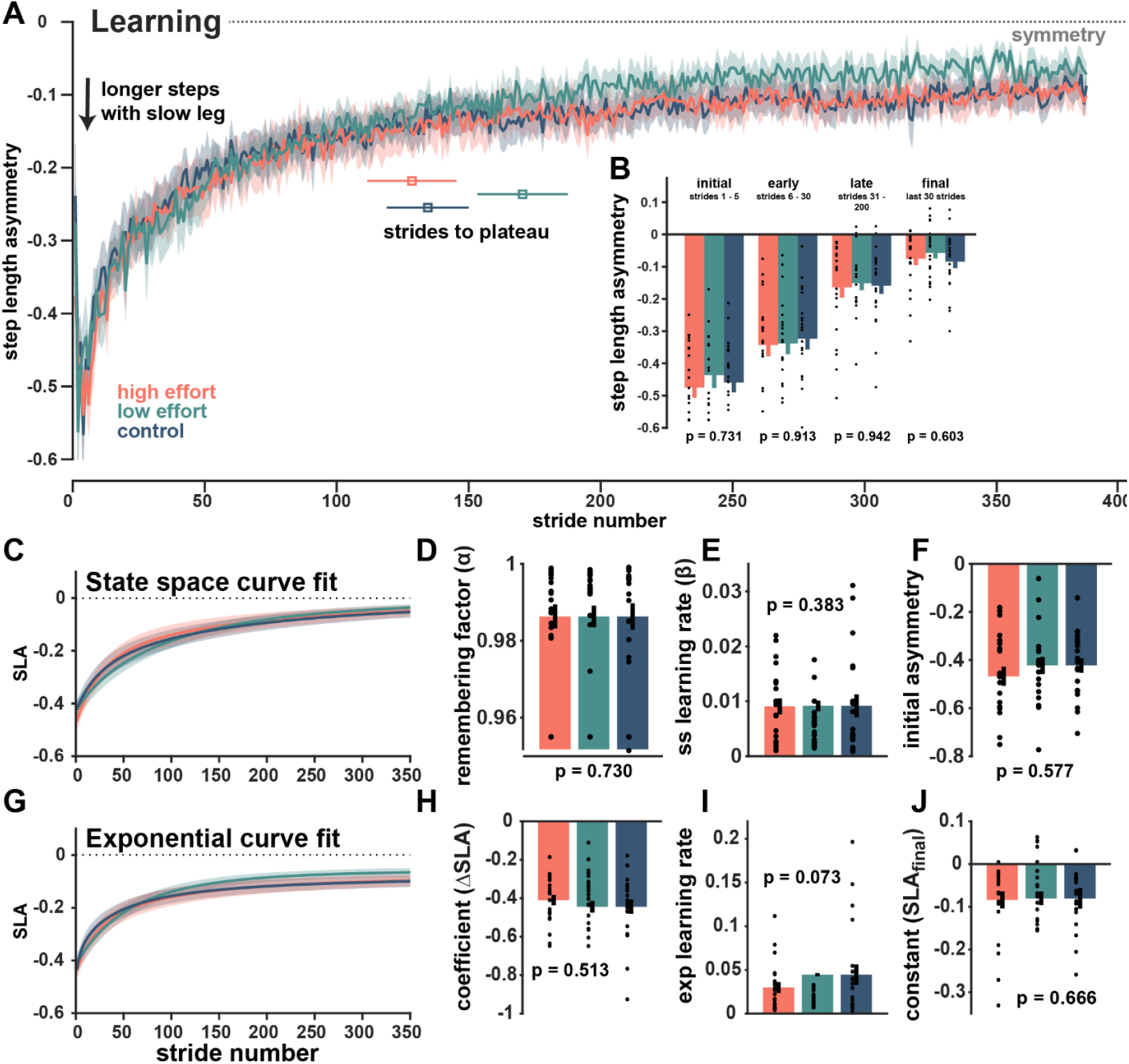
Step length asymmetry during learning. A) Step length asymmetry over strides of the split-belt perturbation. Asymmetry for all three groups progresses from longer steps on the slow leg toward equal length steps. All three groups’ step lengths remained biased toward longer steps on the slow legs at the end of the 10-minute split-belt exposure. Strides to plateau (+/- standard error) are also plotted against the x-axis for each group. B) binned step length asymmetry for initial, early, late, and final periods during the learning block. Each bar represents the mean and standard error across subjects for each bin during learning. C) the average of individually fitted state space learning fits with shaded standard error. D-F) state space model parameters including remembering factor (D), learning rate (E), and the initial asymmetry (F). G) the average of individually fitted exponential curves with shaded standard error. H-J) exponential fit parameters, including the coefficient, representing the change in step length asymmetry from initial to late learning (H), the exponential learning rate (I), and the exponential fit constant, representing the plateau step length asymmetry (J). Dots represent individual participants.

### Increased effort did not influence washout of learning

To assess after affects, we compared aftereffects in a washout block with tied belts at the slow speed (0.5 m/s). We found no differences between effort conditions.

We observed no significant effect of effort condition on step length asymmetry (F(2, 57) = 0.409, p = 0.666), but showed significant changes over the phases of washout, consistent with decaying aftereffects from the split-belt condition (main effect of phase, F(1.48, 84.25) = 183.5, p = 1.50e-27, but no effect of effort condition, p = 0.666, and no interaction, p = 0.706). By the end of the 15-minute washout block, asymmetry was -4.4e-03 ± 4.35e-03 for the high effort group, 0.02 ± 0.004 for the low effort group, and 4.6e-03 ± 2.77e-03 for the control group. When compared against slow baseline, the low effort and control groups had returned to baseline symmetry (low t(18) = 2.07, p = 0.053; control t(19) = 0.47, p = 0.682), but the high effort group remained slightly more asymmetric than baseline (with longer steps on the slow leg) (high t(20) = 2.66, p = 0.015). All three groups were fully symmetric (step length asymmetry equal to zero during the washout plateau: high t(20) = -0.223, p = 0.826; t(18) = 1.26, p = 0.224; control t(19) = 0.356, p = 0.726).

For washout after initial learning, the first stride to the plateau was 99.3 ± 14.59 strides for the high effort condition, 87.2 ± 15.71 strides for the low effort condition, and 95.0 ± 14.90 strides for the control condition. As in the learning block, effort had no significant effect on strides to plateau during the washout block (one-way ANOVA: F(2, 57) = 0.16, p = 0.855).

*Model-based fits*. Remembering factors and learning rates from the state space model fits were similar between groups (all p > 0.3, Figure 6C-F). Additionally, exponential curve fits of step length asymmetry during washout indicated no differences in the magnitude or decay of aftereffects between groups (all p > 0.08, Figure 6G-J).

**Figure 6.**
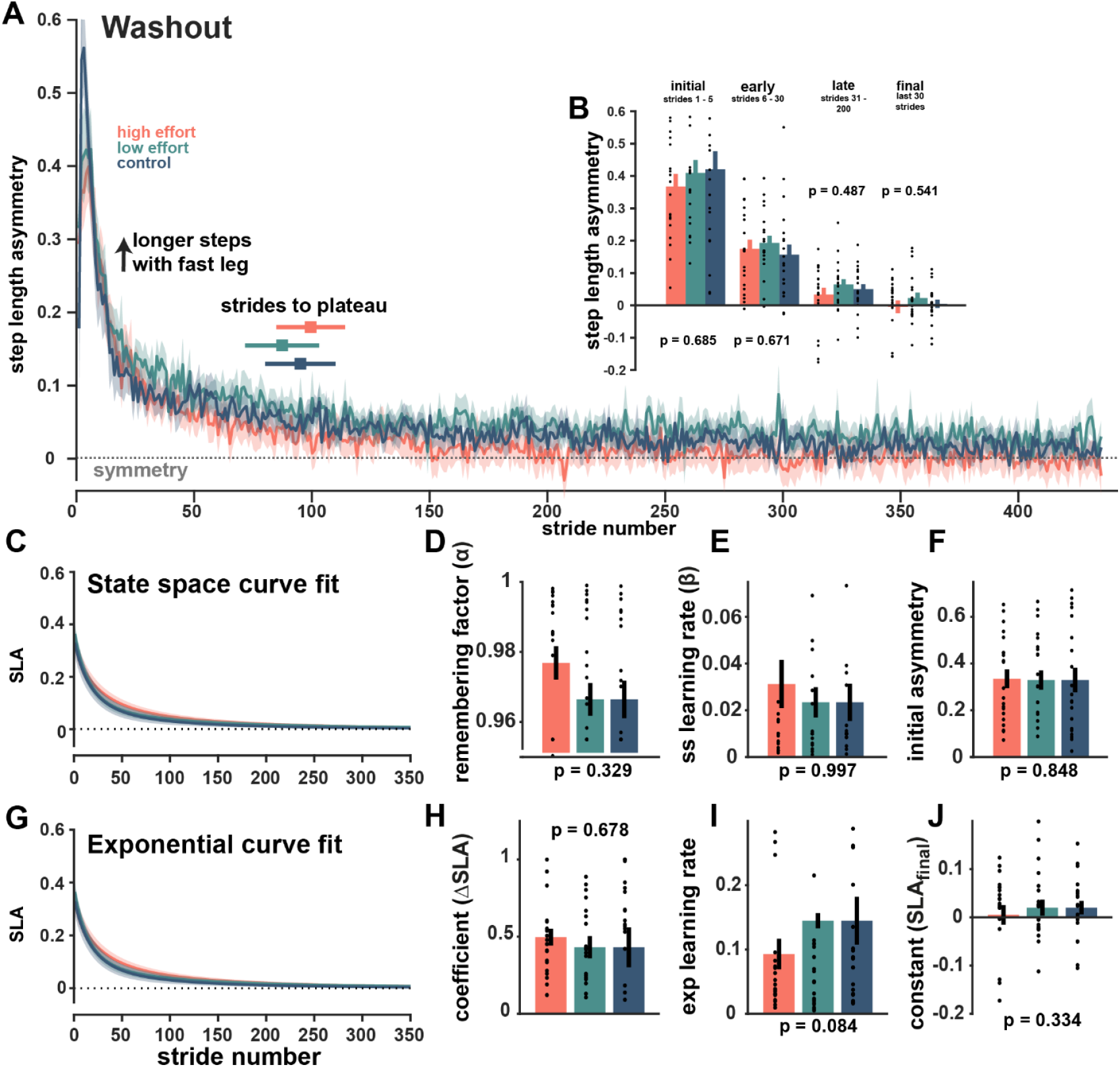
Step length asymmetry during split-belt washout. A) Step length asymmetry over strides of the washout block. Asymmetry for all three groups starts with longer steps on the fast leg, aftereffects from the split-belt perturbation immediately prior, and proceeds toward equal length steps over the 15-minute washout. Strides to plateau (+/- standard error) are also plotted against the x-axis for each group. B) binned step length asymmetry for initial, early, late, and final periods during the washout block. Each bar represents the mean and standard error across subjects for each bin. C) the average of individually fitted state space learning fits with shaded standard error. D-F) state space model parameters including remembering factor (D), learning rate (E), and the initial asymmetry (F). G) the average of individually fitted exponential curves with shaded standard error. H-J) exponential fit parameters, including the coefficient, representing the change in step length asymmetry from initial to late washout (H), the exponential learning rate (I), and the exponential fit constant, representing the plateau step length asymmetry at the end of washout (J). Dots are individual participants.

### Ground reaction force asymmetry adaptation in washout was unaffected by effort condition

Peak propulsive, braking, and vertical ground reaction force asymmetries and propulsive impulse asymmetry were insensitive to effort condition (RM-ANOVA for propulsion F(2,60) = 0.065, p = 0.937; braking, F(2,60) = 1.491, p = 0.233; vertical, F(2,60) = 2.063, p = 0.136; impulse, F(2,60) = 0.410, p = 0.665). All were sensitive to the phase of learning: peak propulsive forces were initially negative but increased toward symmetry (F(1.79,107.5) = 105.9, p = 1.18e-44); peak braking force was initially positive and decayed across phases back toward symmetry (F(1.93,116.0) = 176.5, p = 2.42e-35); vertical ground reaction forces asymmetries were initially positive, but rapidly decayed in the early and later phases (F(1.10, 66.16) = 68.1, p = 2.15e-12). Impulse asymmetries were initially negative and adjusted back toward symmetry (F(1.46, 87.68) = 1.78, p = 1.00e-12). None of the interactions were significant (propulsion F(3.58, 107.5) = 0.897, p = 0.937; braking, F(3.87, 116.0) = 0.121, p = 0.972; vertical, F(2.21,66,16) = 2.072, p = 0.129; impulse, F(2.92, 87.68) = 1.78, p = 0.159). Changing braking, vertical ground reaction force, and impulse asymmetries demonstrated kinetic adaptation, but our task effort manipulation did not create group differences in the adaptation of braking and propulsion asymmetries during washout.

### Effort condition did not affect relearning

In relearning, step length asymmetry differed based on phase, but not based on effort condition (RM_ANOVA, phase, F(1.48, 84.25) = 183.5, p = 1.50e-27, effort condition, F(2,57) = 0.409, p = 0.666, and interaction, F(2.96, 84.25) = 0.463, p = 0.706). Because of visual differences on the relearning asymmetry plot (Figure 7A), we also report the pairwise comparisons between each of the effort groups for each of binned asymmetry phases (Table 1). While there is a trend towards lower asymmetry at Plateau in the low group compared to control, this did not reach significance (p=0.086).

**Table 1:** Pairwise comparisons of step length asymmetry between effort conditions in phases of the relearning block.

|  | Initial | Early | Late | Plateau |
| --- | --- | --- | --- | --- |
| **High v Low** | $p = 0.317$ | $p = 0.440$ | $p = 0.453$ | $p = 0.323$ |
| **High v Control** | $p = 0.740$ | $p = 0.785$ | $p = 0.463$ | $p = 0.481$ |
| **Low v Control** | $p = 0.457$ | $p = 0.252$ | $p = 0.119$ | $p = 0.086$ |

**Figure 7.**
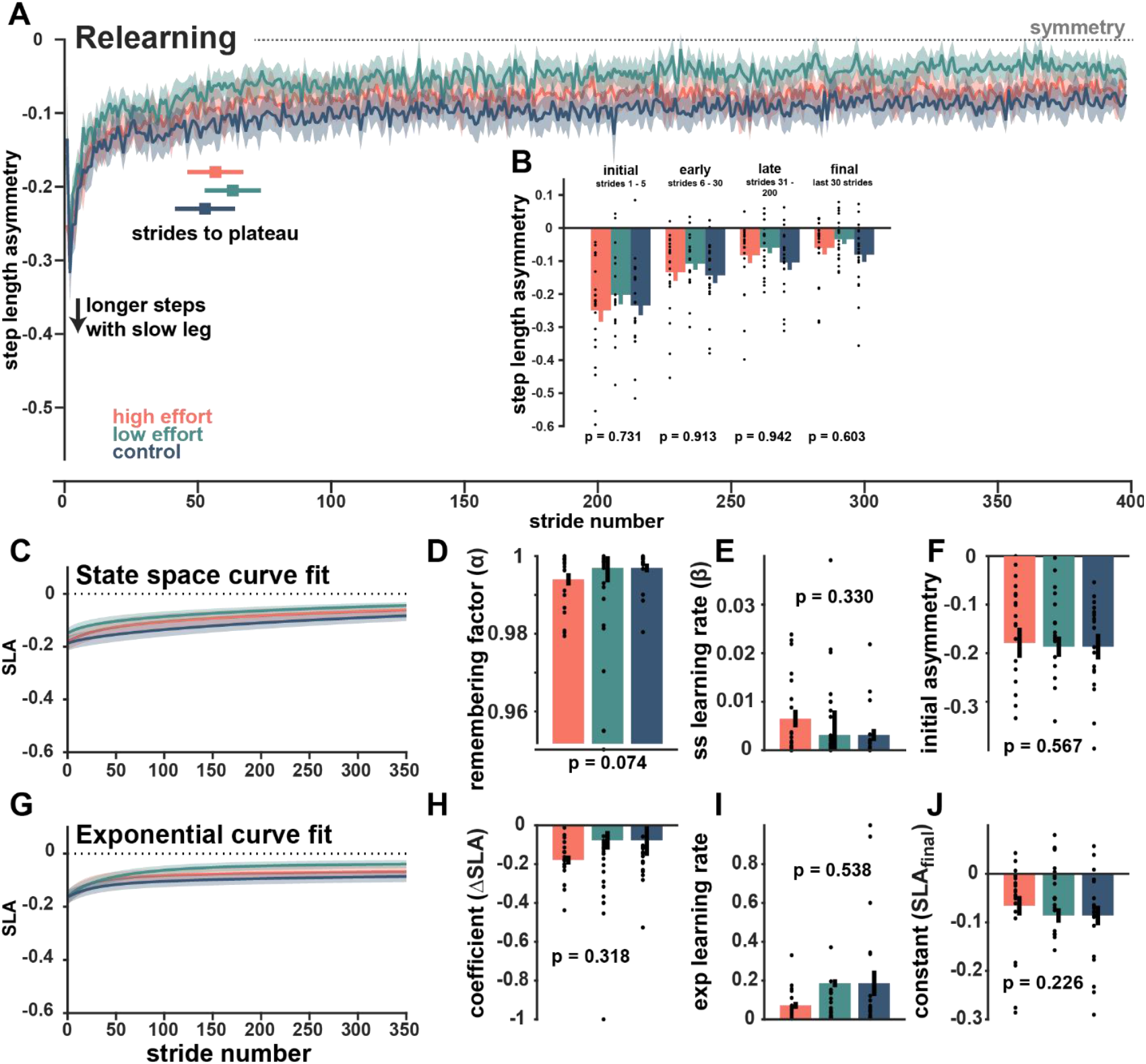
Step length asymmetry during split-belt relearning. A) Step length asymmetry over strides of relearning, the second split-belt perturbation. Asymmetry for all three groups progresses from longer steps on the slow leg toward equal length steps. All three groups’ step lengths remained biased toward longer steps on the slow legs at the end of the 10-minute relearning block. Strides to plateau (+/- standard error) for relearning are also plotted against the x-axis for each group. B) binned step length asymmetry for initial, early, late, and final periods during the relearning block. Each bar represents the mean and standard error across subjects for each bin. C) the average of individually fitted state space learning fits for the relearning block. D-F) state space model parameters including remembering factor (D), learning rate (E), and the initial asymmetry (F). G) the average of individually fitted exponential curves for relearning. H-J) exponential fit parameters, including the coefficient, representing the change in step length asymmetry from initial to late relearning (H), the exponential learning rate (I), and the exponential fit constant, representing the plateau step length asymmetry (J). Dots are individual participants.

At the end of participants’ second split-belt exposure, the step length asymmetry plateau for the high effort group was -0.06 ± 0.005, for the low effort group was -0.03 ± 0.003, and the control effort group was -0.08 ± 0.005. When compared to slow baseline walking, none of the three groups reached full symmetry during the relearning block (paired t-test comparing plateau relearning and slow baseline step length asymmetry: high t(20) = -4.24, p = 3.97e-04; low t(18) = -2.47, p= 0.029; control F(19) -4.54, p = 2.23e-04). Similarly, relearning plateau was asymmetric when compared to perfect symmetry (step length asymmetry = 0) for all groups (high t(20) = -2.89, p= 0.009; low t(18) = -2.23, p= 0.038; control t(19) = -3.76, p = 0.001).

*Model fits*. State space models fits had similar parameters for all groups during relearning (all p > 0.07, Figure 7C-F). Similarly, exponential fits had similar parameters for all groups during relearning (all p > 0.2, Figure 7G-J).

### All groups exhibited savings

We compared step length asymmetry phase measures from learning and relearning split-belt exposures to assess savings. In a three-way ANOVA, step length asymmetry showed a main effect of block (learning or relearning) and phase, with lower asymmetry in the relearning block (phase, F(1.67, 85.12) = 6.73e-35, p = 0.036, and block, F(1, 51) = 384.8, p = 1.32e-36). However, there was no effect of effort condition (effort condition, F(2,51) = 0.453, p = 0.512). Lower asymmetry, especially in the initial and early phases of relearning, is consistent with savings and also explains the significant interaction between phase and block (F(1.76, 89,70) = 89.56, p = 7.12e-49). No other interactions were significant: effort condition and phase, F(3.34, 85.12), 0.217, p = 0.902; effort condition and block F(2, 51) = 0.728, p = 0.488; effort condition with phase and block, F(3.52, 89.70) = 0.325, p = 0.838).

All groups reached a plateau in fewer strides during relearning: 56.6 ± 10.52 strides for the high effort condition (paired ttest between learning and relearning, t(16) = 3.74, p = 1.78e-04), 63.1 ± 10.56 strides for the low effort condition (t(17) = 4.29, p = 4.92e-04), and 52.7 ± 11.31 strides for the control condition (t(18) = 4.06, p = 7.31e-04), but did not differ between groups (one-way ANOVA between effort conditions in relearning, F(2,56) = 0.220, p = 0.807). A repeated measures ANOVA between groups in learning and relearning confirmed a main effect of relearning (F(1,51) = 48.75, p = 5.18e-09) with no effect of effort condition and no interaction (effort condition F(2,51) = 1.37, p = 0.261, interaction F(2, 51) = 0.59, p = 0.560).

### Model-based fits

We assess savings based on the model fit parameters using repeated measures ANOVAs with block (learning or relearning) and effort condition and paired tests between learning and relearning parameters (Table 2). In the state space learning model, savings was revealed in the initial asymmetry values, 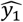, which were lower in relearning for all groups (RM-ANOVA with a significant main effect of block, F(1,51) = 345.9, p = 2.24e-24, and no main effect of or interaction with effort condition, F(2,51) = 0.522, p = 0.597 and F(2,51) = 0.396, p = 0.675). However, there was no effect of block or effort condition on remembering factor or learning rate (remembering factor with no main effect of block, F(1,51) = 2.64, p = 0.111, or effort condition, F(2, 51) = 0.052, p = 0.949, no interaction, F(2,51) = 2.68, p = 0.078; learning rate with no main effect of block, F(1,51) = 0.289, p = 0.593; or effort condition, F(2,51) = 0.557, p = 0.576 ; no interaction F(2,51) = 1.86, p = 0.167).

**Table 2:** Model fit learning metrics compared between learning and savings.

| Comparing learning and relearning (paired t-test between learning and relearning) | High effort group | Low effort group | Control group | ANOVA (effort condition x block) |
| --- | --- | --- | --- | --- |
| State space remembering | $T(17) = -2.42$, $P = 0.028$ | $T(17) = 0.84$, $P = 0.415$ | $T(18) = -1.74$, $P = 0.098$ | Effort condition, $p = 0.949$; block, $p = 0.111$; interaction, $p = 0.078$ |
| States space learning rate | $T(17) = 2.17$, $P = 0.045$ | $T(17) = -1.46$, $P = 0.162$ | $T(18) = 0.42$, $P = 0.677$ | Effort condition, $p = 0.576$; block, $p = 0.593$; interaction, $p = 0.167$ |
| State space initial asymmetry | $P(17) = -10.5$, $P = 1.46e-08$ | $T(17) = -11.1$, $P = 3.20e-09$ | $T(18) = -10.7$, $P = 3.20e-09$ | Effort condition, $p = 0.597$; block, $p = 2.24e-24$; interaction, $p = 0.675$ |
| Exponential coefficient | $T(17) = -4.30$, $P = 0.5.475-04$ | $T(17) = -2.77$, $P = 0.013$ | $T(18) = -3.66$, $P = 0.0.002$ | Effort condition, $p = 0.647$; block, $p = 6.26e-07$; interaction, $p = 0.246$ |
| Exponential learning rate | $T(17) = -1.78$, $P = 0.094$ | $T(17) = -2.21$, $P = 0.041$ | $T(18) = -2.45$, $P = 0.0246$ | Effort condition, $p = 0.374$; block, $p = 4.84e-04$; interaction, $p = 0.726$ |
| Exponential constant | $T(17) = -2.89$, $P = 0.011$ | $T(17) = -3.44$, $P = 0.003$ | $T(18) = -2.46$, $P = 0.024$ | Effort condition, $p = 0.374$; block, $p = 0.4.91e-06$; interaction, $p = 0.488$ |

Exponential fit parameters demonstrated savings for all groups. Exponential coefficient changed based on block (RM-ANOVA with main effect of block, F(1,51) = 32.4, p = 6.26e-07), but no effect of effort condition, (F(2,51) = 0.439, p = 0.647), or interaction (F(2,51) = 1.44, p = 0.246). Exponential learning rates were higher in learning than in relearning (main effect of block F(1,51) = 13.9, p = 4.84e-04), with no effect of or interaction with effort condition (effort condition, F(2, 51) = 0.745, p = 0.480, and no interaction, F(2,51) = 0.322, p = 0.726). Similarly, exponential constants were smaller in relearning (F(1,51) = 26.1, p = 4.91e-06), but with no effect of or interaction with effort conditions (effort condition F(2,51) = 1.00, p = 0.374, interaction F(2,51) = 0.727, p = 0.488).

In summary, step length asymmetry, strides to plateau, state-space model initial asymmetry values, and exponential fit coefficients, learning rates and constants all exhibited savings, with no effect of effort condition.

### Ground reaction force asymmetries also demonstrated savings

Similar to ground reaction forces in learning, peak propulsive, braking, and vertical ground reaction force asymmetries and propulsive impulse asymmetry were insensitive to effort condition (RM-ANOVA for propulsion F(2,60) = 0.749, p = 0.749; braking F(2,60) = 0.716, p = 0.493; vertical F(2,60) = 0.683, p = 0.509; impulse F(2,60) = 0.955, p = 0.391). Peak propulsion asymmetry was unchanged over learning phases (F(1.59, 95.64) = 0.058, p = 0.058), trending toward as slight decrease in positive asymmetry (no interaction F(3.19, 95.64) = 1.64, p = 0.184).

As in learning, peak braking asymmetry was initially biased toward the slow leg and adjusted past symmetry to favor the fast leg with positive asymmetry (phase F(1.63, 97.64) = 318.3, p = 3.97e-40; no interaction F(3.25,97.64) = 0.919, p = 0.441). Peak vertical force was initially negative and increased toward symmetry (F(1.56, 93.44) = 17.65, p = 2.87e-06; no interaction F(3.11, 93.44) = 0.171, p = 0.922). Finally, impulse asymmetry was positive, but decaying over phases (F(1.59, 95.27) = 30.25, p = 1.72e-09; no interaction F(3.18, 95.27) = 1.09, p = 0.360).

Ground reaction force asymmetries also demonstrated savings. Three-way ANOVAs between effort conditions, within phase and block (learning or relearning), showed no effect of effort (propulsion asymmetry, F(2, 60) = 0.796, p = 0.468; braking asymmetry F(2,60) = 0.439, p = 0.647; vertical asymmetry F(2, 60) = 0.561, p = 0.573; impulse asymmetry F(2,60) = 0.709, p = 0.496). Block (learning versus relearning) affected peak propulsive force asymmetry, which was slightly lower on average in relearning (main effect of block F(1,60) = 12.55, p = 0.001, interaction between effort condition and block F(2, 60) = 1.48, p = 0.235; effort condition and phase F(3.63, 108.94) = 0.833, p = 0.428; block and phase F(2.15, 128.7) = 1.66, p = 0.193; three-way interaction F(4.29, 128.7) = 1.44, p = 0.223); ttest between initial force asymmetry in learning and savings, effort conditions combined, initial phase: t(62) = -2.11, p = 0.039, early t(62) = -2.66, p = 0.011, late t(62) = -2.85, p = 0.006).

Block, phase, and the interaction between the two were significant for all other ground reaction force asymmetries, indicating that the changes in each phase differed between the learning and relearning blocks. In peak braking asymmetry, participants started nearer to symmetry (less negative asymmetry) in relearning and remained higher until the plateau phase (main effect for block, F(1, 60) = 382, p = 1.03e-27, and for phase, F(1.96, 118) = 656, p = 4.19e-64, and interaction F(1.72, 103.24) = 85.1, p = 9.09e-21; nonsignificant interaction between effort condition and block F(2,60) = 0.254, p = 0.777; effort condition and phase F(3.92, 117.65) = 1.09, p = 0.364; three way interaction F(3.44, 103.2) = 0.290, p = 0.858; ttest initial t(62) = -10.1, p = 8.85e-15, early t(62) = -21,57, p < 2.2e-16, late t(62) = -14.78, p < 2.2e-16, plateau t(62) = -0.266, p = 0.768).

Peak vertical ground reaction force asymmetry, participants started and remained nearer to symmetry (less negative asymmetry) in relearning than in learning until the plateau phase (RM-ANOVA main effect for block, F(1, 60) = 78.16, p = 1.83e-12, and for phase, F(1.77, 105.9) = 67.56, p = 2.80e-18, and interaction, F(1.64, 98.24) = 8.84, p = 7.15e-04; nonsignificant interactions between effort condition and block F(2, 60) = 0.156, p = 0.856, effort condition and phase F(3.53, 105.9) = 0.370, p = 0.806, three way interaction F(3.27, 98.24) = 0.346, p = 0.809; ttest, initial t(62) = -4.52, p = 2.84e-05, early t(62) = -8.05, p = 3.25e-11, late t(62) = -6.57, p = 1.17e-08, plateau t(62) = -1.29, p = 0.203).

Propulsive impulse asymmetry is positive, but decaying across phases; in relearning, impulse was closer to symmetry in the initial and early phases (RM-ANOVA main effect for block, F(2,60) = 0.709, p = 5.20e-05, and for phase, F(1.78, 107.0) = 35.46, p = 1.08e-11, interaction between phase and block, F(2.24, 134.5) = 9.51, p = 7.09e05, three way interaction between effort condition, block and phase F(4.48, 134.5) = 3.55, p = 0.006; nonsignificant interactions of effort and block F(2, 60) = 2.95, p = 0.600 and effort condition and phase F(3.57, 107.1) = 0.462, p 0.742; ttest, t(62) = 3.02, p = 0.004, t(62) = 4.90, p = 7.10e-06, t(62) = 1.03, p = 0.306, t(62) = 1.28, p = 0.207).

Overall ground reaction force asymmetries indicated savings in the relearning block. Patterns of adaptation in braking and vertical forces and propulsive impulse paralleled observations in learning but demonstrated faster adjustment to the plateau value in relearning.

### Step length was similar between groups during learning, washout, and relearning

While step length asymmetry was similar between groups, we confirmed that the underlying step lengths were also comparable between groups. Step length changed based on leg (consistent with the split-belt perturbation) and phase, but not on the basis of effort condition (RM-ANOVA with main effects of leg, F(1, 58) = 306, p = 8.26e-25, and phase, F(2.24,129.8) = 148.5, p = 2.68e-36, but no effect of effort condition, F(2,58) = 0.097, p = 0.908). The interaction between leg and phase was significant, consistent with adaptation to the split-belt (F(2.05,119.0) = 312.6), p = 1.37e-48). The interaction between effort condition and phase was also significant (F(4.48, 129.8) = 2.41, p = 0.046), but pairwise analysis based on estimated marginal means showed no differences between effort conditions based on the combinations of phase and fast leg. Step lengths showed a similar pattern in washout, changing across phases (F(1.83,104.6) = 89.1, p = 3.07e-22) and varying between legs, with a significant interaction, all features consistent with split-belt aftereffects (leg F(1,57) = 144.2, p = 3.00e-17, and interaction F(1.73, 98.33) = 2.96, p = 5.62e-40). In relearning, step lengths were longer for the slow leg (F(1,58) = 98.7, p = 3.93e-14) and changed over the learning phases (F(1.33, 76.9) = 0.546, p = 1.04e-13) with a significant interaction (leg:phase F(1.47,85.01) = 155.9, p = 2.56e-25).

### Step time and step time asymmetry were similar between groups during learning, washout, and relearning

Comparing step time between groups, fast leg, and across the phases of learning, indicated that step times got progressively longer on the fast leg, but no effect of effort condition (three-way RM-ANOVA with main effects of leg, F(1,58) = 515, p = 1.50e-30, and phase, F(1.87, 108.2) = 159, p = 7.40e-32, but no effect of effort, F(2,58) = 0.349, p = 0.707; leg:phase interaction, F(1.66, 96.22) = 24.77, p = 2.08e-08, other interaction terms were not significant, all p > 0.268). Similarly, in washout and relearning, step times had no main effect of effort condition effect (p > 0.827). In relearning, the three-way interaction between effort condition, phase and leg was significant (F(2.16, 62.6) = 4.23, p = 0.017). Estimated marginal means analysis revealed a trending difference between the control and high effort groups’ fast leg in the initial phase, with longer step times in the high effort group (t(58) = -2.39, p = 0.061, Bonferroni corrected).

Step time asymmetry was similar to previously published data: at the beginning of learning, step time asymmetry was near zero during initial adaptation to split-belt walking, but quickly increased to a positive steady state asymmetry (24). Positive asymmetry indicated longer duration steps on the fast leg (driven by a longer duration of swing). Across learning, washout, and relearning, step time asymmetry had no main effect of effort condition (all p > 0.200), but was sensitive to phase in learning and washout, consistent with step time asymmetry increasing during the learning process and decreasing toward symmetry during washout (both p < 0.007). In relearning, step time asymmetry was recovered so quickly and saved so completely that there was no effect of phase (p = 0.344).

### Step width and step width asymmetry were similar between groups during learning, washout, and relearning

As part of our check for changes in stability, we compared step widths during adaptation blocks. Across learning, washout, and savings, step widths were unaffected by effort condition (all p > 0.186). Step width asymmetry was similarly indifferent to effort condition (all p > 0.255).

### Step variability measures had mixed sensitivity to effort condition and phase during learning

We examined step length and step time variability as a potential proxy for exploration that might drive faster adaptation. We assessed step width variability as a balance measure. We used the same binned learning phases to calculate step metric variance measures across the learning block, then used three-way ANOVAs to look for effects of effort condition, fast leg, and phase within an adaptation block. Step length variability was unaffected by effort condition across learning, washout, and relearning (all F(2,58) < 2.46, all p > 0.245). Similarly, step time variability was similar between effort groups across adaptation blocks (F(2,57) <1.28, all p > 0.285).

Step width variability was sensitive to effort condition in the learning block, with higher variability in the control group especially in the initial and early blocks (main effect of effort condition F(2,58) = 3.34, p = 0.043, trending interaction between effort condition and phase F(2.05, 59.49) = 3.06, p = 0.053). This effect did not persist in washout and relearning (both F(2,5) < 2.09, p > 0.133). However, in relearning the interaction between effort condition and leg (F(2,58) = 3.68, p = 0.031) as well as the interaction between effort condition, leg, and phase were significant F(2, 58.06) = 3.44, p = 0.039). Assessing estimated marginal means, we compared the three effort groups for each combination of phase and fast leg. The only significant comparison was between the high effort and control effort groups in the initial phase for the slow leg, which showed higher variability in the control group (t(58) = 3.01, p = 0.012, Bonferroni corrected). Because the initial phase contains only 5 strides, differences in variability are not likely to be indicative true differences between the two groups.

In all adaptation blocks, step width variability decreased after the initial phase as they became familiar with the block (learning F(1.03,59.49) = 253, p = 2.36e-23; washout F(1.04,59.1) = 272, p = 3.65e-24; relearning F(1.02, 58.95) = 230, p = 3.71e-22).

In summary, additional kinematic measures of gait such as step length, step width, step time asymmetry, and their variance, all revealed no significant effect of effort condition.

### Among participants who completed both high and low effort conditions, savings was strong

A subset of participants in the high and low effort groups returned to the lab on a separate day to complete the walking task in the opposite effort condition. During the second visit, we compared the high-low and low-high groups on their third and fourth exposures to the split-belt perturbation and examined savings across both visits.

Across the learning, washout, and relearning blocks, there were no significant differences between the high-low and low-high groups. Comparing step length asymmetry across binned phases, the main effects for effort condition were F(1,18) < 2.14, and p > 0.161 for all three blocks. State space and exponential model fits were also similar with t(11) > |1.79| and p > 0.101. Strides to plateau were similar (t(15.9) = 0.563, p = 0.581).

Within the second visit, comparisons between exposure 3 (visit 2; learning) to exposure 4 (visit2; relearning), revealed small same day improvements, with significant improvements in the state space initial value (t(9) = - 3.12, p = 0.012), exponential constant t(9) = -2.38, p = 0.041). All other comparisons were unchanged between second visit learning and relearning (all p > 0.08).

We used a three-way ANOVA to assess the phases of learning and relearning across both visits. There was no main effect of effort group (low-high and high-low, F(1,15) = 0.347, p = 0.565). In all split-belt exposure blocks, participants adjusted closer to symmetry across phases (main effect of phase F(1.54, 23.14) = 80.17, p = 2.56e-10). Across exposures, participants also got closer to symmetry (main effect of exposure F(1.89,28.35) = 74.05, p = 9.12e-12) and did so in earlier phases (interaction between phase and exposure F(3.44, 51.67) = 24.03, p = 1.58e-10). No other interactions were significant (all p > 0.645).

Thus, once again, we see learning and savings in all groups irrespective of the effort condition in the prior session. Importantly, there is no significant effect of effort condition on the rate or extent of learning.

**Figure 8:**
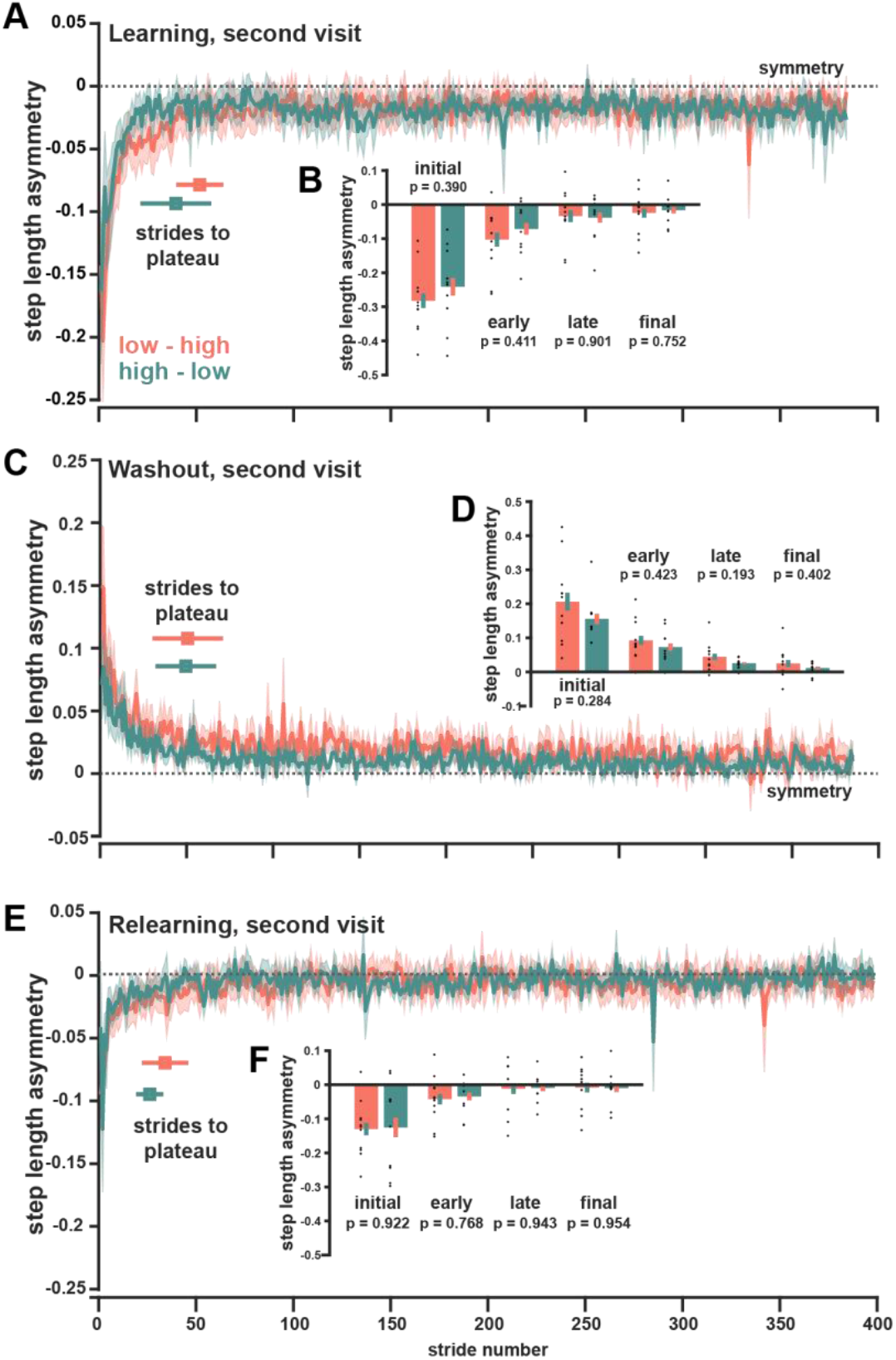
Step length asymmetry during learning (A), washout (C), and relearning (E) for the second visit in the opposite effort condition. Nested subplots are included showing the binned step length asymmetry for initial, early, late, and plateau bins during learning (B), washout (D), and relearning (F) from the second visit. Throughout the figure, the color indicates the current (second visit) effort condition. Shaded regions and error bars are ±1SEM. Dots are individual participants.

## DISCUSSION

Effort cost reduction observed during learning poses the question of whether the motor system is trying to reduce error and effort in parallel? Which cost is more important in driving or shaping the learning process? We compared learning in split-belt locomotion between three groups with differing levels of background effort. We found no differences between high, low, and control effort conditions in the rate or extent of step length asymmetry adaptation. The three groups also had similar aftereffects and similar patterns of relearning. A subset of individuals returned to complete the task in the opposite effort condition. Second visit participants in both groups showed substantial savings between the two visits with no effect of the effort condition in which they first encountered the split-belt perturbation.

Based on prior work related to effort, movement, and learning, we could rationalize three potential ways in which effort might interact with motor learning. In the first, we anticipated accelerated learning because the higher magnitude effort during early learning might increase the urgency of error reduction. We considered that effort corollaries like increased neuromuscular recruitment or augmented proprioceptive feedback might increase implicit salience of the effort gradient or that higher percieved effort costs might more readily engage explicit learning mechanisms. Previous work has shown faster visuomotor rotation learning with effortful muscle co-contraction, which could be attributed to greater effort landscape sensitivity (51). Work in locomotion has shown that disrupted proprioception delayed learning, we considered that the effect may also work in the opposite direction (38). In the second possibility, we examined effort’s potential to impair learning. We posited potential mechanisms like distraction, which has been shown to slow learning in locomotion and reaching (18, 52). Alternatively, effort costs may reduce the “goodness” of the learning environment, interacting with tonic dopamine in the brain, and reducing motivation (44). While both rationales are compelling, our findings do not point towards either an accelerative or slowing effect of effort on learning.

Our results align most closely with our final possibility: effort might not affect the general rate or extent of step length asymmetry adaptation. Previous work looking at responses to effort landscapes have indicated perseveration in their preferred movement patterns even when they are no longer optimal. In some cases, force exploration of the environment can that people prompt motor adjustments, such as gait adjustments to reduce exoskeleton torque or adopt positive step length asymmetry after being guided through the potential effort landscape (9, 10). In other studies, people persist in their preferred movement patterns despite experience in more favorable conditions. In force field reaching, participants maintain an effortful, straight-line reaching path even after being taught a low effort curved path (54). In locomotion, people persist in their preferred step frequency in a cost landscape defined by inspired ratios of CO_2_ and O_2_ (55). Even after free and forced exploration across a range of step frequencies where low CO_2_ - high O_2_ gas mixes were available for non-preferred frequencies, people chose to maintain their preferred cadence in a high CO_2_ - low O_2_ condition despite higher percieved exertion (55). The range of these predictions for effort’s role in motor learning reflect the general ambiguity of our understanding of how effort and error interact with movement and motor learning; our study provides evidence that error signals and a gradual response to them may supersede any response to an effort gradient.

*How much effort is enough?* Since faster learning was observed in split-belt locomotion on an incline, we compared the relative effort costs of the two protocols (29).The incline task increased propulsion, decreased braking and could not delineate whether elevated effort was contributing to accelerated learning and provides a useful point of comparison for “how much” effort might be sufficient to enhance learning. Based on data from Butterfield and Collins, the metabolic cost of spilt-belt walking is similar to the cost of tied belt walking at the average belt speed (32). Their data includes a small correction for belt-speed difference, but since both our study and the incline study set belt speeds to 0.5 and 1.5 m/s we neglected this correction in order to apply tied-belt walking cost equations for added mass and incline (32). Ludlow and Weyland investigated the net metabolic cost of walking relating incline, added mass, and walking speed (56). In agreement with other findings, they found that the costs incurred with additional mass were proportional to the mass as a percentage of body mass (33, 56). Based on an their empirical equation, we estimated the net metabolic cost of walking to be 2.61 W/kgBM with the 15% body mass and 6.76 W/kgBM on an 8.5 degree incline (29, 56). The same equation predicts a gross metabolic cost of 2.27 W/kgBM for level, unweighted walking at 1.0 m/s, so that the additional mass constituted a proportional 15% increase in net metabolic cost, while the incline increased the cost by nearly 200%. Thus, if the faster learning observed in the inclined split-belt is attributable to higher effort magnitude, effort costs from any reasonable added mass may be insufficient to drive faster learning.

In selecting the added mass, we wanted to choose a meaningful effort cost without accumulating fatigue over the course of the session. Oxygen consumption is significantly higher than unweighted walking with as little as 10% body mass of additional mass and masses below 25% of body mass are manageable for sustained periods (33, 57). The high effort group, carrying 15% BM of additional mass fell comfortably in this range. Based on early and late learning net metabolic costs reported by Finley et al. (2013), we assume that the added mass generates a uniformly proportional increase in net metabolic cost during learning, so participants in the high effort group stand to reduce net metabolic cost by nearly 20% during the learning block (from roughly 4.37 W/kgBM in early learning to roughly 3.51 W/kgBM in late learning based on multiplying reported early and late split-belt cost values by 1.15; (16). Notably, the magnitude of this decrease in metabolic cost nears previous reports for the average just noticeable difference (58). In other locomotion tasks people highly are sensitive even to small differences in effort and can adjust their gait to new optima implicitly (10, 11). When wearing an exoskeleton that penalized preferred step frequency, participants adjusted away from their preferred gait for cost savings of 4-8%, though cost gradient alone was insufficient to initiate adaptation and participants needed to be guided to explore the environment before adjusting away from their preferred gait (10, 59). In a study that allowed participants to select a treadmill belt speed by changing their step placement asymmetry, participants *generated* asymmetry to walk at a faster speed (symmetric walking set the treadmill to 0.5 m/s) which minimized cost of transport (60). Though we observed no effect of effort on learning, these studies suggest a high-resolution sensitivity to changes in metabolic cost and imply that i) high effort group experienced meaningfully more effort during the walking task and ii) the difference in effort cost created by the additional weight is sufficient to incentivize gait changes.

We considered several reasons that effort might have no effect on split-belt learning, aftereffects, or relearning. Since we modulated the effort necessary for the split-belt walking task by adding extra mass, a familiar experience in day-to-day walking, it is possible that the increased effort was low salience in the context of split-belt walking and its inherent discomfort, instability, and novelty. Since the background effort was inherent to the split-belt task (which was novel to every participant), perhaps the effort had no effect because the effort landscape was entirely foreign. Alternatively, effort might affect learning in ways that are not apparent in step length asymmetry. In forcefield reaching, pain had mixed effects on the progress of error reduction, but consistently changed motor learning strategies, including muscle recruitment and feedforward preparation (3, 43). In asymmetric walking, pain had no effect on adaptation to visually cued, gradually increasing step length asymmetry, but impaired 24-hour retention (61). Among participants that visited the lab a second time, we saw no evidence of an effect of effort cost on retention, but an effect may have been diluted by the switching between effort conditions. Fatigue similarly changes patterns of muscle recruitment and, in some tasks, increases error during learning and recall (62, 63). These findings attest to the resilience of the learning process, which can compensate for pain and fatigue without compromising error reduction (43, 61). Effort may similarly induce subtle changes in the patterns or strategies of movements without changing learning and our data further suggests that aftereffects and relearning are also unaffected by background effort.

Since effort did not impair motor learning, modest increases in effort have clinical potential as a tool to build strength and endurance in rehabilitation without slowing motor recovery. This may offer insight into findings that post-stroke patients gait training at higher intensity showed greater improvements in clinical outcomes like the 6-minute walk test and comfortable gait speed than did post-stroke patients training at low intensity (64, 65).

*Limitations*. We specifically selected the high effort mass to increase metabolic cost without accumulating fatigue, for this reason the metabolic costs are not equivalently comparable to some other locomotor learning tasks. Thus, our experimental design cannot rule out an effect of a higher magnitude of effort.

Because participants wore the vest for the entire duration of the split-belt task, they completed adaptation, washout, and readaptation in the same motor context. Further research will need to address whether effort cost experienced during learning might generalize to unweighted walking conditions and whether the extent or generalization is sensitive to effort during learning. Other than instructing participants to look ahead and not at the belts, we also made no adjustments to correct for the proprioceptive mismatch of treadmill walking where there is no optical flow. While this is similar to most split-belt studies, removing vision during adaptation improves learning by mitigating the visual-proprioceptive mismatch (31). Since effort might increase proprioceptive gain, learning might be more sensitive to effort-related modulations when proprioceptive information is not countered by stationary visual feedback. Future work could explore effort and adaptation with obscured vision or in virtual reality environments with optical flow. This study included only healthy participants and thus cannot directly indicate the effect of different effort costs on a clinical population who will face different obstacles and effort landscapes during gait retraining. We did not collect rate of percieved exertion. Doing so would have helped determine the extent to which participants in the two groups experienced the effort costs as different. Collecting rate of percieved exertion between each experimental block could have also helped us determine how subject effort evolved over the experiment and whether that evolution differed between effort conditions.

Many prior split-belt studies identified moderate to extremely large effect sizes: f-test effect sizes that we calculated from reported data or estimated from published figures ranged from 0.266 to 3.68. These findings were reported from studies with 16 and 8 participants per group respectively, and our groups sizes were consistent with most published split-belt walking data (18, 22, 29, 66–68). Based on our survey of the literature, metrics reported as key results, groups of 9 - 17 participants usually provided sufficient power. We selected the primary studies for our power analysis because we hypothesized that attention and pain mechanisms might be similar to those of effort. We have tried to collect data from a moderate number of participants to achieve adequate statistical power to detect effect sizes similar to these split-belt studies. Based on a statistical power of 0.95 with a 0.05 significance level, our one-way ANOVAs between three groups should be sensitive to effect sizes greater than *f =* 0.507. We also conducted pair-wise t-tests between groups, these tests should be sensitive to effects sizes greater than *d =* 1.03. Though similar to effect sizes observed in other studies, both of these thresholds are classified as large effects, and our study may fail to identify moderate or small effects of effort. Thus, our results showed an absence of any *large* effect of background effort on motor learning; however, the effect of effort may be more subtle than previously published effects and may fall below the sensitivity we were able to achieve with one-way ANOVAs for three experimental groups and with t-tests between groups (18, 29, 61).

## Conclusion

This study investigated the effect of background effort on split belt adaptation, retention, and readaptation during one or two visits to the laboratory. Using step length asymmetry as our learning metric, we identified no effect on rate or extent of adaptation, rate or extent of readaptation, or on retention within or across visits. The absence of any effect of background effort on the learning process suggests that task effort costs do not change the perception of the belt speeds, urgency of error or effort reduction, or the use of explicit strategy during split-belt adaptation. These findings could imply that modest increases in task effort do not impair learning and so may be beneficial for building strength during rehabilitation or motor retraining.

## REFERENCES

1. Galea JM, Mallia E, Rothwell J, Diedrichsen J. The dissociable effects of punishment and reward on motor learning. Nat Neurosci 18:597–602, 2015. doi: 10.1038/nn.3956.

2. Nikooyan AA, Ahmed AA. Reward feedback accelerates motor learning. Journal of Neurophysiology 113:633–646, 2015. doi: 10.1152/jn.00032.2014.

3. Salomoni SE, Marinovic W, Carroll TJ, Hodges PW. Motor Strategies Learned during Pain Are Sustained upon Pain-free Reexposure to Task. Medicine & Science in Sports & Exercise 51:2334, 2019. doi: 10.1249/MSS.0000000000002059.

4. Takahashi CD, Nemet D, Rose-Gottron CM, Larson JK, Cooper DM, Reinkensmeyer DJ. Effect of muscle fatigue on internal model formation and retention during reaching with the arm. J Appl Physiol (1985) 100:695–706, 2006. doi: 10.1152/japplphysiol.00140.2005.

5. Carlisle RE, Kuo AD. Optimization of energy and time predicts dynamic speeds for human walking. eLife 12:e81939, 2023. doi: 10.7554/eLife.81939.

6. Donelan JM, Kram R, Arthur D. K. Mechanical and metabolic determinants of the preferred step width in human walking. Proc R Soc Lond B 268:1985–1992, 2001. doi: 10.1098/rspb.2001.1761.

7. Ralston HJ. Energy-speed relation and optimal speed during level walking. Int Z Angew Physiol Einschl Arbeitsphysiol 17:277–283, 1958. doi: 10.1007/BF00698754.

8. Sánchez N, Park S, Finley JM. Evidence of Energetic Optimization during Adaptation Differs for Metabolic, Mechanical, and Perceptual Estimates of Energetic Cost. Sci Rep 7:7682, 2017. doi: 10.1038/s41598-017-08147-y.

9. Sánchez N, Simha SN, Donelan JM, Finley JM. Taking advantage of external mechanical work to reduce metabolic cost: the mechanics and energetics of split-belt treadmill walking. J Physiol 597:4053–4068, 2019. doi: 10.1113/JP277725.

10. Selinger JC, O’Connor SM, Wong JD, Donelan JM. Humans Can Continuously Optimize Energetic Cost during Walking. Current Biology 25:2452–2456, 2015. doi: 10.1016/j.cub.2015.08.016.

11. McAllister MJ, Blair RL, Donelan JM, Selinger JC. Energy optimization during walking involves implicit processing. Journal of Experimental Biology 224:jeb242655, 2021. doi: 10.1242/jeb.242655.

12. Shadmehr R, Huang HJ, Ahmed AA. Effort, reward, and vigor in decision-making and motor control. Curr Biol 26:1929–1934, 2016. doi: 10.1016/j.cub.2016.05.065.

13. Summerside EM, Shadmehr R, Ahmed AA. Vigor of reaching movements: reward discounts the cost of effort. Journal of Neurophysiology 119:2347–2357, 2018. doi: 10.1152/jn.00872.2017.

14. Huang HJ, Kram R, Ahmed AA. Reduction of Metabolic Cost during Motor Learning of Arm Reaching Dynamics. J Neurosci 32:2182–2190, 2012. doi: 10.1523/JNEUROSCI.4003-11.2012.

15. Huang HJ, Ahmed AA. Reductions in muscle coactivation and metabolic cost during visuomotor adaptation. Journal of Neurophysiology 112:2264–2274, 2014. doi: 10.1152/jn.00014.2014.

16. Finley JM, Bastian AJ, Gottschall JS. Learning to be economical: the energy cost of walking tracks motor adaptation. The Journal of Physiology 591:1081–1095, 2013. doi: 10.1113/jphysiol.2012.245506.

17. Gonzalez-Rubio M, Velasquez NF, Torres-Oviedo G. Explicit Control of Step Timing During Split-Belt Walking Reveals Interdependent Recalibration of Movements in Space and Time [Online]. Frontiers in Human Neuroscience 13, 2019. https://www.frontiersin.org/articles/10.3389/fnhum.2019.00207 [9 Feb. 2023].

18. Malone LA, Bastian AJ. Thinking About Walking: Effects of Conscious Correction Versus Distraction on Locomotor Adaptation. Journal of Neurophysiology 103:1954–1962, 2010. doi: 10.1152/jn.00832.2009.

19. Mawase F, Haizler T, Bar-Haim S, Karniel A. Kinetic adaptation during locomotion on a split-belt treadmill. Journal of Neurophysiology 109:2216–2227, 2013. doi: 10.1152/jn.00938.2012.

20. Reisman DS, Wityk R, Silver K, Bastian AJ. Locomotor adaptation on a split-belt treadmill can improve walking symmetry post-stroke. Brain 130:1861–1872, 2007. doi: 10.1093/brain/awm035.

21. Day KA, Leech KA, Roemmich RT, Bastian AJ. Accelerating locomotor savings in learning: compressing four training days to one. Journal of Neurophysiology 119:2100–2113, 2018. doi: 10.1152/jn.00903.2017.

22. Leech KA, Roemmich RT, Bastian AJ. Creating flexible motor memories in human walking. Sci Rep 8:94, 2018. doi: 10.1038/s41598-017-18538-w.

23. Reisman DS, Block HJ, Bastian AJ. Interlimb Coordination During Locomotion: What Can be Adapted and Stored? Journal of Neurophysiology 94:2403–2415, 2005. doi: 10.1152/jn.00089.2005.

24. Stenum J, Choi JT. Step time asymmetry but not step length asymmetry is adapted to optimize energy cost of split-belt treadmill walking. The Journal of Physiology 598:4063–4078, 2020. doi: 10.1113/JP279195.

25. Malone LA, Vasudevan EVL, Bastian AJ. Motor Adaptation Training for Faster Relearning. Journal of Neuroscience 31:15136–15143, 2011. doi: 10.1523/JNEUROSCI.1367-11.2011.

26. Mariscal DM, Iturralde P, Torres-Oviedo G. Altering attention to split-belt walking increases the generalization of motor memories across walking contexts. Journal of neurophysiology null: null, 2020. doi: 10.1152/jn.00509.2019.

27. Ogawa T, Kawashima N, Ogata T, Nakazawa K. Predictive control of ankle stiffness at heel contact is a key element of locomotor adaptation during split-belt treadmill walking in humans. J Neurophysiol 111:722–732, 2014. doi: 10.1152/jn.00497.2012.

28. Roemmich RT, Long AW, Bastian AJ. Seeing the errors you feel enhances locomotor performance but not learning. Curr Biol 26:2707–2716, 2016. doi: 10.1016/j.cub.2016.08.012.

29. Sombric CJ, Calvert JS, Torres-Oviedo G. Large Propulsion Demands Increase Locomotor Adaptation at the Expense of Step Length Symmetry [Online]. Frontiers in Physiology 10, 2019. https://www.frontiersin.org/articles/10.3389/fphys.2019.00060 [25 Jul. 2023].

30. Song CN, Stenum J, Leech KA, Keller CK, Roemmich RT. Unilateral step training can drive faster learning of novel gait patterns. Sci Rep 10:18628, 2020. doi: 10.1038/s41598-020-75839-3.

31. Torres-Oviedo G, Bastian A. Seeing Is Believing: Effects of Visual Contextual Cues on Learning and Transfer of Locomotor Adaptation. The Journal of Neuroscience 30:17015– 17022, 2010. doi: 10.1523/JNEUROSCI.4205-10.2010.

32. Butterfield JK, Collins SH. The energy cost of split-belt walking for a variety of belt speed combinations. Journal of Biomechanics 132:110905, 2022. doi: 10.1016/j.jbiomech.2021.110905.

33. Bastien GJ, Willems PA, Schepens B, Heglund NC. Effect of load and speed on the energetic cost of human walking. Eur J Appl Physiol 94:76–83, 2005. doi: 10.1007/s00421-004-1286-z.

34. Huang TP, Kuo AD. Mechanics and energetics of load carriage during human walking. Journal of Experimental Biology 217:605–613, 2014. doi: 10.1242/jeb.091587.

35. Wei K, Körding K. Uncertainty of feedback and state estimation determines the speed of motor adaptation [Online]. Frontiers in Computational Neuroscience 4, 2010. https://www.frontiersin.org/article/10.3389/fncom.2010.00011 [10 Jun. 2022].

36. Hoogkamer W, Bruijn SM, Potocanac Z, Van Calenbergh F, Swinnen SP, Duysens J. Gait asymmetry during early split-belt walking is related to perception of belt speed difference. Journal of Neurophysiology 114:1705–1712, 2015. doi: 10.1152/jn.00937.2014.

37. Yokoyama H, Sato K, Ogawa T, Yamamoto S-I, Nakazawa K, Kawashima N. Characteristics of the gait adaptation process due to split-belt treadmill walking under a wide range of right-left speed ratios in humans. PLoS ONE 13:e0194875, 2018. doi: 10.1371/journal.pone.0194875.

38. Hubbuch JE, Bennett BW, Dean JC. Proprioceptive feedback contributes to the adaptation toward an economical gait pattern. Journal of Biomechanics 48:2925–2931, 2015. doi: 10.1016/j.jbiomech.2015.04.024.

39. Mariscal DM, Iturralde PA, Torres-Oviedo G. Altering attention to split-belt walking increases the generalization of motor memories across walking contexts. Journal of Neurophysiology 123:1838–1848, 2020. doi: 10.1152/jn.00509.2019.

40. Bouffard J, Bouyer LJ, Roy J-S, Mercier C. Tonic Pain Experienced during Locomotor Training Impairs Retention Despite Normal Performance during Acquisition. J Neurosci 34:9190–9195, 2014. doi: 10.1523/JNEUROSCI.5303-13.2014.

41. Dancey E, Murphy B, Andrew D, Yielder P. Interactive effect of acute pain and motor learning acquisition on sensorimotor integration and motor learning outcomes. Journal of Neurophysiology 116:2210–2220, 2016. doi: 10.1152/jn.00337.2016.

42. Galgiani JE, French MA, Morton SM. Acute pain impairs retention of locomotor learning. Journal of Neurophysiology 131:678–688, 2024. doi: 10.1152/jn.00343.2023.

43. Lamothe M, Roy J-S, Bouffard J, Gagné M, Bouyer LJ, Mercier C. Effect of Tonic Pain on Motor Acquisition and Retention while Learning to Reach in a Force Field. PLOS ONE 9:e99159, 2014. doi: 10.1371/journal.pone.0099159.

44. Niv Y, Daw ND, Joel D, Dayan P. Tonic dopamine : opportunity costs and the control of response vigor. Dopamine - revisited 191:507–520, 2007.

45. Buurke TJW, Lamoth CJC, van der Woude LHV, den Otter R. Handrail Holding During Treadmill Walking Reduces Locomotor Learning in Able-Bodied Persons. IEEE Transactions on Neural Systems and Rehabilitation Engineering 27:1753–1759, 2019. doi: 10.1109/TNSRE.2019.2935242.

46. Herzfeld DJ, Shadmehr R. Motor variability is not noise, but grist for the learning mill. Nat Neurosci 17:149–150, 2014. doi: 10.1038/nn.3633.

47. McAndrew Young PM, Dingwell JB. Voluntarily changing step length or step width affects dynamic stability of human walking. Gait & Posture 35:472–477, 2012. doi: 10.1016/j.gaitpost.2011.11.010.

48. Owings TM, Grabiner MD. Variability of step kinematics in young and older adults. Gait & Posture 20:26–29, 2004. doi: 10.1016/S0966-6362(03)00088-2.

49. Torres-Oviedo G, Vasudevan E, Malone L, Bastian AJ. Locomotor adaptation. In: Progress in Brain Research. Elsevier, p. 65–74.

50. Sombric CJ, Torres-Oviedo G. Augmenting propulsion demands during split-belt walking increases locomotor adaptation of asymmetric step lengths. J NeuroEngineering Rehabil 17:69, 2020. doi: 10.1186/s12984-020-00698-y.

51. Heald JB, Franklin DW, Wolpert DM. Increasing muscle co-contraction speeds up internal model acquisition during dynamic motor learning. Sci Rep 8:16355, 2018. doi: 10.1038/s41598-018-34737-5.

52. Taylor JA, Thoroughman KA. Divided Attention Impairs Human Motor Adaptation But Not Feedback Control. Journal of Neurophysiology 98:317–326, 2007. doi: 10.1152/jn.01070.2006.

53. Galgiani JE, French MA, Morton SM. Acute pain impairs retention of locomotor learning. Journal of Neurophysiology 131:678–688, 2024. doi: 10.1152/jn.00343.2023.

54. Kistemaker DA, Wong JD, Gribble PL. The Central Nervous System Does Not Minimize Energy Cost in Arm Movements. Journal of Neurophysiology 104:2985–2994, 2010. doi: 10.1152/jn.00483.2010.

55. Wong JD, O’Connor SM, Selinger JC, Donelan JM. Contribution of blood oxygen and carbon dioxide sensing to the energetic optimization of human walking. Journal of Neurophysiology 118:1425–1433, 2017. doi: 10.1152/jn.00195.2017.

56. Ludlow LW, Weyand PG. Estimating loaded, inclined walking energetics: No functional difference between added and body mass. In: 2016 IEEE 13th International Conference on Wearable and Implantable Body Sensor Networks (BSN). 2016 IEEE 13th International Conference on Wearable and Implantable Body Sensor Networks (BSN), p. 306–311.

57. Puthoff ML, Darter BJ, Nielsen DH, Yack HJ. The Effect of Weighted Vest Walking on Metabolic Responses and Ground Reaction Forces. Medicine & Science in Sports & Exercise 38:746, 2006. doi: 10.1249/01.mss.0000210198.79705.19.

58. Medrano RL, Thomas GC, Rouse EJ. Can humans perceive the metabolic benefit provided by augmentative exoskeletons? J NeuroEngineering Rehabil 19:26, 2022. doi: 10.1186/s12984-022-01002-w.

59. Simha SN, Wong JD, Selinger JC, Abram SJ, Donelan JM. Increasing the gradient of energetic cost does not initiate adaptation in human walking. Journal of Neurophysiology 126:440–450, 2021. doi: 10.1152/jn.00311.2020.

60. Roemmich RT, Leech KA, Gonzalez AJ, Bastian AJ. Trading symmetry for energy cost during walking in healthy adults and persons post-stroke. Neurorehabil Neural Repair 33:602–613, 2019. doi: 10.1177/1545968319855028.

61. Galgiani JE, French MA, Morton SM. Acute pain impairs retention of locomotor learning. Journal of Neurophysiology 131:678–688, 2024. doi: 10.1152/jn.00343.2023.

62. Branscheidt M, Kassavetis P, Anaya M, Rogers D, Huang HD, Lindquist MA, Celnik P. Fatigue induces long-lasting detrimental changes in motor-skill learning. eLife 8:e40578, 2019. doi: 10.7554/eLife.40578.

63. Enoka RM. Neuromechanics of Human Movement. Human Kinetics, 2015.

64. Boyne P, Miller A, Kubalak O, Mink C, Reisman DS, Fulk G. Moderate to Vigorous Intensity Locomotor Training After Stroke: A Systematic Review and Meta-analysis of Mean Effects and Response Variability. Journal of Neurologic Physical Therapy 48:15, 2024. doi: 10.1097/NPT.0000000000000456.

65. Holleran CL, Rodriguez KS, Echauz A, Leech KA, Hornby TG. Potential Contributions of Training Intensity on Locomotor Performance in Individuals With Chronic Stroke. Journal of Neurologic Physical Therapy 39:95, 2015. doi: 10.1097/NPT.0000000000000077.

66. French MA, Morton SM, Charalambous CC, Reisman DS. A locomotor learning paradigm using distorted visual feedback elicits strategic learning. Journal of Neurophysiology 120:1923–1931, 2018. doi: 10.1152/jn.00252.2018.

67. Mawase F, Shmuelof L, Bar-Haim S, Karniel A. Savings in locomotor adaptation explained by changes in learning parameters following initial adaptation. Journal of Neurophysiology 111:1444–1454, 2014. doi: 10.1152/jn.00734.2013.

68. Vasudevan EVL, Bastian AJ. Split-Belt Treadmill Adaptation Shows Different Functional Networks for Fast and Slow Human Walking. Journal of Neurophysiology 103:183–191, 2010. doi: 10.1152/jn.00501.2009.

